# Restraint of TGFβ family signaling by SMAD7 is necessary for hematopoietic stem cell maturation in the embryo

**DOI:** 10.1101/2025.08.23.671940

**Authors:** Laura F. Bennett, Wenbao Yu, Chia-Hui Chen, Hyun Hyung An, Joanna Tober, Kai Tan, Nancy A. Speck

## Abstract

Hematopoietic stem cells (HSCs), defined as cells that can engraft an adult when transplanted, mature from precursors (pre-HSCs) that differentiate from hemogenic endothelial cells (HECs) in the embryo. Many signaling pathways required to generate the first hematopoietic stem and progenitor cells in the embryo are well-characterized, but how HSCs mature from pre-HSCs is poorly understood. Here we show that “mothers against decapentaplegic homolog 7” (SMAD7), a negative regulator of transforming growth factor beta (TGFβ) and bone morphogenetic protein (BMP) signaling, is required for pre-HSC to HSC maturation. Deletion of *Smad7* in endothelial cells allows the formation of pre-HSCs from HECs but impairs their maturation into HSCs. The data indicate that although TGFβ and BMP signaling are required for the generation of HECs and for HECs to undergo an endothelial-to-hematopoietic transition to generate pre-HSCs, one or both pathways must be subsequently down-regulated for effective pre-HSC to HSC maturation.

## Introduction

Hematopoietic ontogeny in the mouse embryo occurs in the yolk sac and in the major caudal arteries (dorsal aorta, vitelline, and umbilical) in several waves, producing hematopoietic stem and progenitor cells (HSPCs) with different lineage potentials^1^. Most HSPCs, except yolk sac-derived primitive erythroid and erythroid/megakaryocytic progenitors, differentiate from hemogenic endothelial cells (HECs) through an endothelial-to-hematopoietic transition (EHT) during which HECs round up, lose their tight junctions, and disengage from the endothelium^2^. Newly formed HSPCs in the caudal arteries accumulate in intra-arterial hematopoietic clusters (IAHCs) that remain briefly attached to the luminal endothelium before being released into the circulation to colonize secondary hematopoietic organs such as the fetal liver and thymus.

Functional assays, *in vivo* fate mapping, and barcoding experiments in mice identified multiple types of HSPCs produced from HECs in the yolk sac and caudal arteries, including hematopoietic stem cells (HSCs), their immediate precursors (pre-HSCs), and HSC-independent progenitors that are not derived from HSCs^1,3^. HSC-independent progenitors include erythro-myeloid progenitors (EMPs) in the yolk sac ^4,5^, and lympho-myeloid biased progenitors (LMPs) in the yolk sac and caudal arteries. LMPs primarily give rise to B and T lymphocytes and myeloid cells in culture, but are not capable of either directly engrafting or maturing *ex vivo* into an HSC ^1,3,6-12^. *In vivo*, LMPs contribute to the first wave of thymus-seeding progenitors (TSPs), which colonize the mouse thymus beginning at embryonic day (E) 11.25 and produce invariant γδ T cells and lymphoid tissue inducer cells (LTi), preparing the thymus for a second influx of adult-like, likely HSC-derived TSPs several days later ^13-16^. HSC-independent HSPCs in the caudal arteries also include multipotent progenitors (MPPs) that produce a more balanced repertoire of erythroid, megakaryocyte, myeloid, and lymphoid cells in *ex vivo* cultures, but similar to LMPs, cannot engraft^17,18^. More recent in vivo barcoding experiments identified embryonic MPPs (eMPPs) in E10.5 embryos, before HSC emergence, that contribute to adult hematopoiesis, particularly lymphopoiesis, for at least a year after birth, and are clonally unrelated to HSCs^19^.

HSCs that can directly engraft myelo-ablated adult mice are produced from the caudal arteries, primarily in the aorta-gonad-mesonephros (AGM) region ^20^, and are detectable beginning at E10.5-E11.5^21-23^. HSCs differentiate from precursors called pre-HSCs, which are defined experimentally as cells that cannot directly engraft adult mice but can be matured *ex vivo* into multi-lineage repopulating HSCs^24,25^. It is thought that pre-HSCs and HSC-independent HSPCs differentiate from distinct populations of HECs, with pre-HSCs differentiating from CXCR4^+^ HECs and LMPs/MPPS from CXCR4^-^ HECs in the caudal arteries^17^. Several genes or pathways have been shown to regulate the balance between pre-HSCs/HSCs and HSC-independent HSPCs in the mouse embryo, including Myd88-dependent toll-like receptor signaling, which favors the production of LMPs in the caudal arteries, and the transcription factor EVI1, encoded by the *Mecom* gene, which promotes the formation of HSCs^26,27^.

Compared to the more extensive characterization of pre-HSC and LMP formation, relatively few studies have focused on pre-HSC to HSC maturation. Understanding this process has practical implications, as both efficient pre-HSC generation *and* maturation will be necessary to produce enough transplantable HSCs from human induced pluripotent stem cells (iPSCs) for clinical applications. Pre-HSC to HSC maturation *in vivo* begins in the IAHCs, as transplantation experiments determined that ∼1 adult-repopulating HSC is present in the E11.5 AGM^20,22^. Most pre-HSC to HSC maturation occurs in the fetal liver, which by E12.5 contains ∼50-70 HSCs, very similar to the number of pre-HSCs in the AGM one day prior^20,28,29^. Pre-HSC to HSC maturation is accompanied by the down-regulation of genes and markers involved in embryonic development and vasculogenesis, and up-regulation of genes involved in hematopoietic organ development, lymphoid development, and immune responses^30,31^. Functional markers specific for HSCs, and not present on pre-HSCs, include major histocompatibility complex I (MHC-1) antigens, which are thought to prevent rejection of embryonic HSCs by natural killer cells in adult transplant recipients^28^. Pre-HSC to HSC maturation in IAHCs is accompanied by de-accelerated cell cycling, whereas committed progenitors (likely from the HSC-independent wave) actively cycle ^32^. To our knowledge, no transcription factors or signaling pathways were shown to be specifically required for the maturation of pre-HSCs into HSCs.

We previously performed single-cell transcriptional profiling of cells in the EHT trajectory spanning endothelial cells (ECs) to IAHCs in the caudal arteries and compared the transcriptomes of LMPs and pre-HSCs^10^. We identified ∼900 genes more highly expressed in LMPs and ∼1000 transcripts more abundant in pre-HSCs. Genes more highly expressed in pre-HSCs included several encoding known markers of pre-HSCs and HSCs, such as *Procr, Tnfrf17,* and *Mecom*^27,33,34^, while LMPs expressed higher levels of lympho-myeloid specific genes^10^. *Smad7*, which encodes an inhibitor of transforming growth factor beta (TGFβ) family signaling, was among the most highly differentially expressed genes in pre-HSCs.

SMAD7 inhibits signal transmission from TGFβ family receptors. Canonical TGFβ signaling is triggered by the binding of a TGFβ or bone morphogenetic protein (BMP) ligand to a type II receptor, which phosphorylates and activates its associated type I receptor, which in turn phosphorylates the receptor-regulated R-SMAD transcription factors. The activated R-SMADs associate with the common partner SMAD4, and the complex enters the nucleus as a transcription factor. SMAD7 inhibits TGFβ signaling by disrupting the interaction between the type I receptors and R-SMADs, and by facilitating degradation of the receptors^35^. TGFβ signaling in HECs cooperates with RUNX1 to regulate the expression of genes required for EHT, but the abundance of phosphorylated R-SMADs subsequently decreases in IAHC cells^36-39^. Since *Smad7* expression is elevated in pre-HSCs relative to LMPs, we hypothesized that restraint of TGFβ and/or BMP signaling by SMAD7 may be necessary for generating pre-HSCs or HSCs. We tested this hypothesis by deleting *Smad7* in HECs before EHT, and later in pre-HSCs. We show that SMAD7 is specifically required for the maturation of HSCs from pre-HSCs and is not necessary for producing normal numbers of pre-HSCs or HSC-independent HSPCs.

## Results

### *Smad7* deletion in endothelial cells does not affect the number of IAHC cells but decreases the number of HSCs

We deleted *Smad7* in endothelial cells and examined the formation of HSCs and HSC-independent HSPCs. We used a floxed *Smad7* allele in conjunction with an endothelial-specific, tamoxifen-inducible Cre recombinase (Cdh5-Cre^ERT^)^40^. We injected pregnant dams with tamoxifen at E9.5, before the peak of EHT and IAHC formation from HECs in the caudal arteries. Colony assays of dissociated E10.5 yolk sacs showed no change in the number of EMPs or the distribution between different EMP colony types in *Smad7^f/f^*; Cdh5-Cre^ERT^ (*Smad7*^Δ*/*Δ^) embryos compared to control embryos (Fig. S1A). PCR genotyping of individual colonies from methylcellulose cultures demonstrated that 67% of EMPs contained two deleted *Smad7* alleles (*del/del*), indicating that SMAD7 does not regulate the formation of EMPs (Fig. S1B).

The vasculature of E10.5 *Smad7*^Δ*/*Δ^ embryos was normal as assessed by whole-mount confocal microscopy and visual inspection (Fig. 1A), and the average width of the dorsal aorta was unaltered (Fig. 1B), indicating that any observed effects of SMAD7 loss on HSC and HSPC formation are unlikely to be a secondary consequence of vascular defects. We quantified HECs and the number of IAHC cells in the dorsal aorta within the AGM region at E10.5, when IAHCs are most abundant ^41^. We identified HECs as flat CD31^+^ RUNX1^+^ cells within the wall of the dorsal aorta (RUNX1 is a specific marker of HECs and IAHCs)^42^, and IAHCs as round CD31^+^RUNX1^+^ cells inside the lumen of the vessel^43^ (Fig. 1C, Fig. S1C). We found no alterations in the number of HECs or IAHC cells in *Smad7*^Δ*/*Δ^ embryos (Fig. 1D-E). However, the median fluorescence intensity of RUNX1 was higher in HECs and IAHC cells than in controls (Fig. 1F-G), indicating that SMAD7 deletion had altered their molecular properties, possibly through unrestrained TGFβ and/or BMP signaling or other integrated signaling pathways^35^.

**Figure 1.**
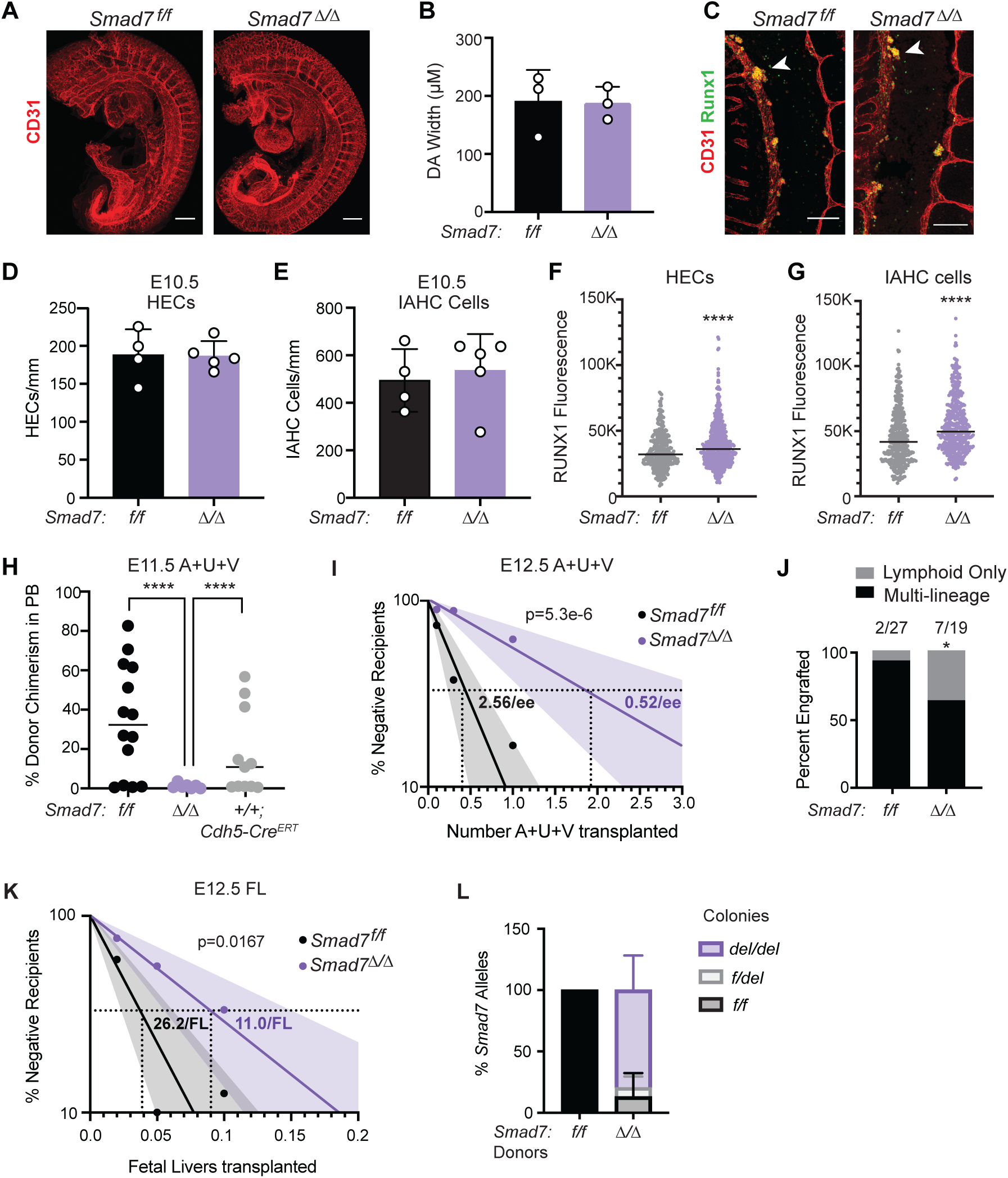
Loss of SMAD7 does not impact IAHC formation but severely reduces the number of adult multi-lineage repopulating HSCs in the caudal arteries. (A) Representative confocal Z-projections of E10.5 *Smad7^f/f^* and *Smad7*^Δ*/*Δ^ embryos (head, intestines, and body wall removed) stained for CD31 (red). Scale bar=250µM. (B) The width of the dorsal aorta (µM) in E10.5 embryos was assessed by taking 5 measurements over 6 sp centered around the vitelline artery. Mean±SD, unpaired, two-tailed Student’s t-test, n=3 embryos per genotype, 29-35 somite pairs (sp). (C) Representative confocal images of IAHC cells (indicated by white arrowheads) in the dorsal aorta of E10.5 embryos stained for CD31 (red) and RUNX (green). Scale bar=100µM, n=4-5 embryos. (D) Quantification of HECs per mm in E10.5 dorsal aortas. Mean±SD, unpaired, two-tailed Student’s t-test. n=4-5 embryos, 29-35 sp. (E) Quantification of IAHC cells in E10.5 dorsal aortas. Mean±SD, unpaired, two-tailed Student’s t-test, n=4-5 embryos. (F,G) Corrected fluorescent intensity of RUNX1 measured from confocal images in HECs (F) or IAHC cells (G) of E10.5 embryos. Mean±SD, unpaired, two-tailed Student’s t-test. n=352 *Smad7^f/f^*HECs, 532 *Smad7*^Δ*/*Δ^ HECs, 364 *Smad7^f/f^* IAHC cells, 425 *Smad7*^Δ*/*Δ^ IAHC cells from 3-4 embryos per genotype. (H) Percent donor-derived CD45^+^ cells in PB of myelo-ablated adult recipient mice 16 weeks after transplanting one embryo equivalent (ee) of E11.5 A+U+V from *Smad7 ^f/f^*, *Smad7*^Δ*/*Δ^, or *Smad7^+/+^*; Cdh5-Cre^ERT^ embryos. Each dot represents one recipient. One-way ANOVA, Tukey’s test for multiple correction comparison, n=7-14 recipients. (I) Limiting dilution analysis of adult-repopulating HSCs in the A+U+V of E12.5 *Smad7^f/f^* and *Smad7*^Δ*/*Δ^ embryos. Shaded regions indicate 95% confidence intervals. HSC frequencies and significant differences in frequencies were determined by ELDA^80^. Data are from 12-21 transplant recipients per cell dose per genotype. Data are plotted as the percentage of recipients at each dose with no donor engraftment. (J) Fraction of engrafted recipients at all doses of donor cells with multi-lineage versus lymphoid-only donor cell contribution. Multi-lineage engraftment is defined as >1% donor contribution to B, T, and myeloid lineages in the PB. Lymphoid-only engraftment refers to recipients with >1% B and/or T chimerism and <1% donor chimerism in myeloid cells in the PB. Fisher’s exact test, two-sided, 95% confidence interval, n=19-27 recipients per genotype. (K) Limiting dilution analysis of adult-repopulating HSCs in the FL of E12.5 *Smad7^f/f^* and *Smad7*^Δ*/*Δ^ embryos. (L) Percentage of allelic deletion in donor-derived LSK cells sorted from the BM of mice transplanted with E12.5 *Smad7^f/f^* and *Smad7*^Δ*/*Δ^ FL cells 16 weeks post-transplant. Donor-derived LSK cells were plated in methylcellulose, and individual colonies were genotyped by PCR. n=17-30 individual colonies. For all panels, *p ≤ 0.05, **p ≤ 0.01, ***p ≤ 0.001, ****p ≤ 0.0001.

We transplanted one embryo equivalent (ee) of AGM regions, umbilical and vitelline arteries (A+U+V) from individual E11.5 embryos into irradiated adult mice to determine if adult repopulating HSCs were affected by SMAD7 loss. No donor-derived cells were observed in the peripheral blood (PB) of recipients transplanted with cells from the A+U+V from E11.5 *Smad7*^Δ*/*Δ^ embryos at 16 weeks post-transplant, whereas 16/25 recipients transplanted with either A+U+V from *Smad7^f/f^* or Cdh5-Cre^ERT^ control embryos exhibited multi-lineage engraftment (Fig. 1H). Notably, there was no evidence of Cre^ERT^ toxicity as there was no difference in engraftment of control cells from *Smad7^f/f^* and Cdh5-Cre^ERT^ embryos (Fig. 1H); therefore, we used *Smad7^f/f^*embryos as controls in most other experiments to maximize breeding efficiency.

Since the frequency of adult repopulating HSCs in the A+U+V of E11.5 *Smad7*^Δ*/*Δ^ embryos was too low to measure, we performed a limiting dilution transplant at E12.5, when there are more HSCs in these sites (approximately 2-3 at E12.5 versus 1 HSC at E11.5) ^22^. Cells from E12.5 *Smad7*^Δ*/*Δ^ A+U+V engrafted, but there was an ∼80% decrease in the number of multi-lineage reconstituting HSCs per embryo, indicating that loss of SMAD7 in endothelial cells severely decreases the number of HSCs in the A+U+V (Fig. 1I). An increased fraction of recipients of E12.5 *Smad7*^Δ*/*Δ^ A+U+V cells also had donor-derived lymphoid cells only in their PB (Fig. 1J, S2A). BM from primary transplant recipients with multi-lineage reconstitution by *Smad7*^Δ*/*Δ^ A+U+V cells contained donor-derived phenotypic LT-HSCs (Lin^-^Sca1^+^c-Kit^+^CD48^-^CD150^+^) (Fig. S2B, C) and could engraft secondary recipient mice (not shown). In contrast, BM from recipients with lymphoid-only reconstitution by *Smad7*^Δ*/*Δ^ A+U+V cells lacked donor-derived LT-HSCs (Fig. S2C) and could not engraft secondary recipient mice. Lymphoid-only reconstituted recipients did, however, contain a small percentage of donor-derived Lin^-^ cells, suggesting that they had been repopulated with a long-lived progenitor with lymphoid potential (Fig. S2D).

The E12.5 fetal liver (FL) contains an estimated 65-70 HSCs due to the migration, expansion, and maturation of pre-HSCs from the AGM region ^20^. We assessed whether the severe reduction in HSCs in the AGM region of *Smad7*^Δ*/*Δ^ embryos could be due to the precocious migration of pre-HSCs from the AGM to the FL by performing a limiting dilution transplant. There was a similar reduction in the estimated number of HSCs per FL in E12.5 *Smad7*^Δ*/*Δ^ embryos (∼60%) compared to the A+U+V (80%) (Fig. 1K). To confirm that recipients were engrafted by LT-HSCs with completely deleted *Smad7* alleles, we sorted donor-derived LSK cells from recipient bone marrows 16 weeks after transplant, plated them in methylcellulose, and genotyped individual colonies. Approximately 80% of sorted donor LSKs from E12.5 FL transplants had bi-allelic excision of *Smad7* (del/del) (Fig. 1L). The data indicate that, although reduced in number in the embryo, SMAD7-deficient HSCs are capable of engrafting adult mice. The data also suggest that premature mobilization to the FL is not responsible for reducing HSCs in the E11.5 and E12.5 AGM regions. In summary, deletion of *Smad7* in endothelial cells did not affect the number of IAHC cells but substantially reduced the number of multi-lineage adult-repopulating HSCs in both the caudal arteries of E11.5-E12.5 embryos and the fetal liver of E12.5 embryos, and it increased the proportion of mice engrafted with long-lived lymphoid-restricted progenitors.

### SMAD7 is not required to generate normal numbers of HSC-independent HSPCs

We determined whether *Smad7* deletion at E9.5 in HECs altered the numbers of HSC-independent progenitors with lymphoid potential in the IAHCs (LMPs), the thymus (TSPs), and in the fetal liver (lympho-myeloid-primed progenitors, or LMPPs). To enumerate LMPs we deleted *Smad7* in endothelial cells at E9.5 and plated limiting numbers of dissociated cells from the AGM regions of E10.5 *Smad7*^Δ*/*Δ^ and control embryos on OP9 stromal cells in the presence of cytokines that promote B and myeloid (M) cell development, then determined the frequency of wells containing B, M, or B+M cells in the cultures 7-10 days later (Fig. 2A). *Smad7*^Δ*/*Δ^ embryos had a similar number of LMPs in their AGM regions as control embryos (Fig. 2B). To confirm *Smad7* was efficiently deleted in LMPs, we flushed the circulating blood containing contaminating yolk sac-derived EMPs from the aortas of dissected AGM regions^44^ before disaggregating the tissue containing the IAHCs, plated the cells in methylcellulose, and PCR genotyped the *Smad7* alleles in individual myeloid colonies. Most (66%) of myeloid colonies had biallelic *Smad7* deletion (Fig. 2C), confirming that SMAD7 is not required for LMP formation. We next determined whether SMAD7 is required for normal numbers of TSPs by quantifying RUNX1^+^ hematopoietic cells within or contacting the keratin-8^+^ thymus primordium in E11.5 *Smad7*^Δ*/*Δ^ and control embryos by confocal microscopy^13^. Deletion of *Smad7* in endothelial cells at E9.5 did not affect the number of RUNX1^+^ TSPs, indicating that SMAD7 is not required for the first wave of TSPs (Fig. 2D, E). Finally, we examined the percentages of phenotypic fetal liver LMPPs (c-Kit^+^CD45^+^Lin^-^ CD135^+^IL7Ra^+^), which are thought to colonize the thymus^13,45,46^, and found no difference between E12.5 *Smad7^Δ/Δ^* and control embryos (Fig. 2F-G). Therefore, by three independent criteria, we established that SMAD7 is not required for generating normal numbers of HSC-independent progenitors with lymphoid potential, including LMPs within the caudal arteries, the first wave of TSPs, and fetal liver LMPPs.

**Figure 2.**
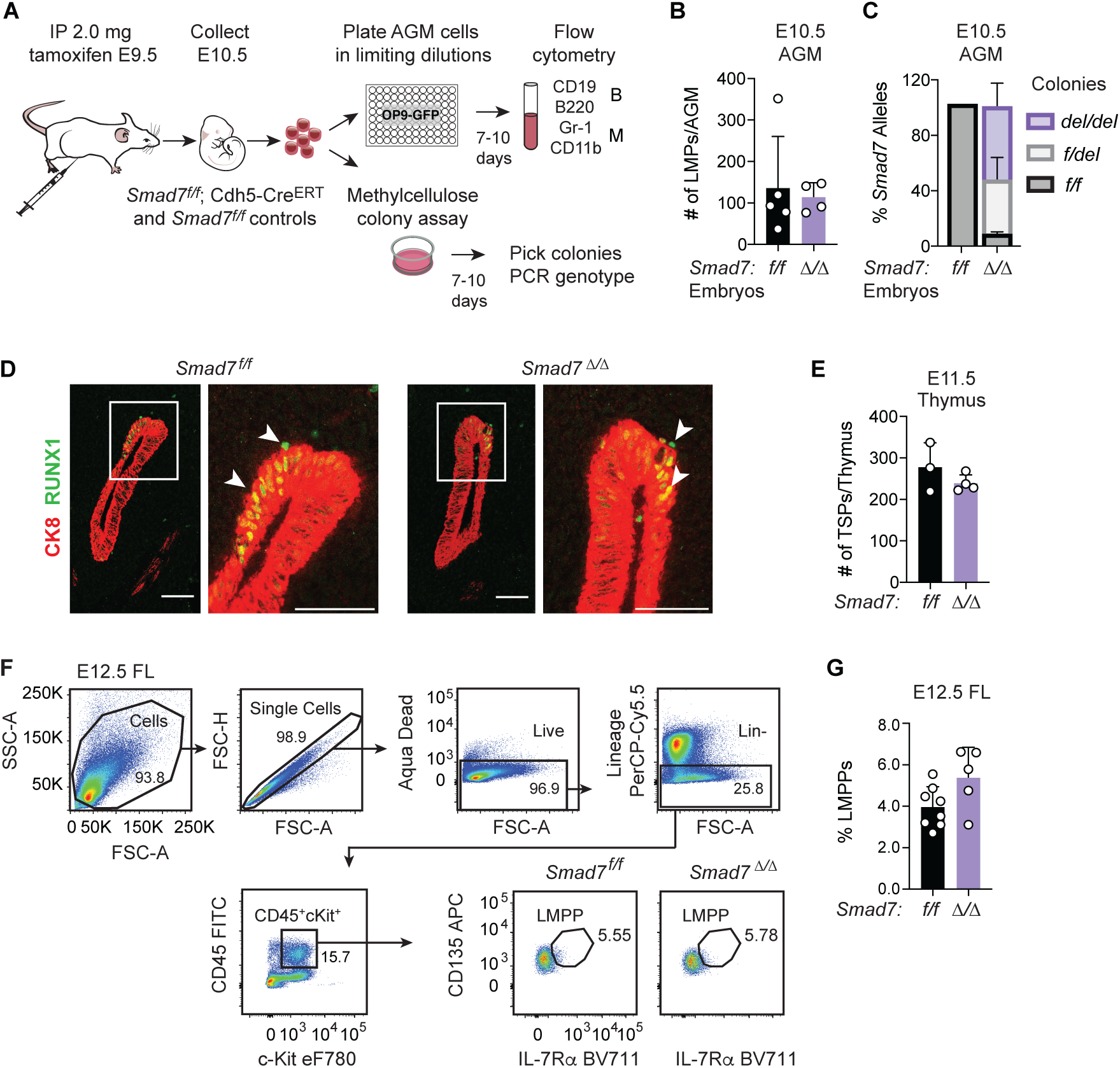
SMAD7 is not required for generating normal numbers of HSC-independent HSPCs. (A) Schematic diagram of the assay for determining the number of LMPs in the AGM regions of E10.5 *Smad7^f/f^* and *Smad7*^Δ*/*Δ^ embryos. Cells from dissociated AGM regions were plated in limiting numbers on OP9 stromal cells, and wells were assessed 7-10 days later for the presence of B and myeloid cells. Cells were also plated in methylcellulose, and individual colonies were picked, and analyzed by PCR to quantify the efficiency of *Smad7* deletion. (B) The number of progenitors per ee that produced B and/or myeloid cells in culture was determined by ELDA^80^. All three colony types (B, B+myeloid, myeloid) are combined in each graph. n=4-5 embryos. (C) Deletion frequency of *Smad7* alleles in myeloid colonies from methylcellulose colony assays. *del/del*, biallelic *Smad7* deletion; *f/del*, monoallelic *Smad7* deletion; *f/f*, no deletion of *Smad7*. n=3 embryos per genotype with 48 individual colonies analyzed per embryo. (D) Representative confocal Z projections of the developing thymus at E11.5 in *Smad7^f/f^* and *Smad7*^Δ*/*Δ^ embryos stained for cytokeratin 8 (CK8) (red) and RUNX1 (green). Scale bar= 150 µM. Enlarged images of the insets (white boxes) are shown to the right of each panel. White arrowheads indicate RUNX1^+^ cells infiltrating or in contact with the thymus. (E) Quantification of RUNX1^+^ TSPs within or contacting the thymus, n=3-4 embryos. (F) Representative flow plots of c-Kit^+^CD45^+^Lin^-^ CD135^+^IL-7Rɑ^+^ LMPPs in E12.5 fetal livers. (G) Percent of LMPPs in Kit^+^CD45^+^Lin^-^ cells in E12.5 fetal livers of *Smad7^f/f^* and *Smad7*^Δ*/*Δ^ embryos. n=5-8 embryos. In all panels mean ± SD, Student’s t-test, unpaired, two-tailed.

### SMAD7 decreases the number of functional pre-HSCs

To determine if the effect of SMAD7 loss on HSCs is mirrored by a decrease in the number of functional pre-HSCs, we performed a limiting dilution pre-HSC analysis. We plated limiting numbers of purified IAHC cells (Ter119^-^CD41^mid/lo/-^CD31^+^CD144^+^ESAM^+^Kit^+^) (Fig. 3A, S3) from E11.5 *Smad7^f/f^*; Cdh5-Cre^ERT^ and control embryos on endothelial cells expressing an activated form of Akt (Akt-ECs), which were previously shown to support pre-HSC to HSC maturation *ex vivo*^47^. We analyzed half of the cells in each well after 4 days for hematopoietic cell growth and for the presence of phenotypic LT-HSCs by flow cytometry and transplanted the remaining cells from each well into lethally irradiated recipient mice to measure the frequency of pre-HSCs that had matured into functional LT-HSCs (Fig. 3A). Hadland *et al.*^48^ demonstrated that LT-HSCs matured from pre-HSCs *ex vivo* are CD45^+^CD144^-^Gr-1^-^F4/80^-^Sca-1^hi^CD201^hi^. All wells exhibited hematopoietic cell growth at all plating densities of both control and *Smad7*^Δ*/*Δ^ IAHC cells (Fig. 3C). However, a smaller fraction of wells seeded with *Smad7*^Δ*/*Δ^ IAHC cells contained phenotypic LT-HSCs at the end of the culture period compared with control cells (Fig. 3B, D). Accordingly, extreme limiting dilution analysis (ELDA) revealed an 85% reduction in functional *Smad7*^Δ*/*Δ^ pre-HSCs capable of maturing into LT-HSCs that engrafted adult recipient mice (Fig. 3E). The 85% decrease in functional pre-HSCs at E11.5 was equivalent to the 80% decrease in LT-HSCs in the E12.5 caudal arteries detected by direct transplantation (Fig. 1I), suggesting that a reduction of functional pre-HSCs causes the decreased number of LT-HSCs in the IAHCs of Smad7^Δ*/*Δ^ embryos.

**Figure 3.**
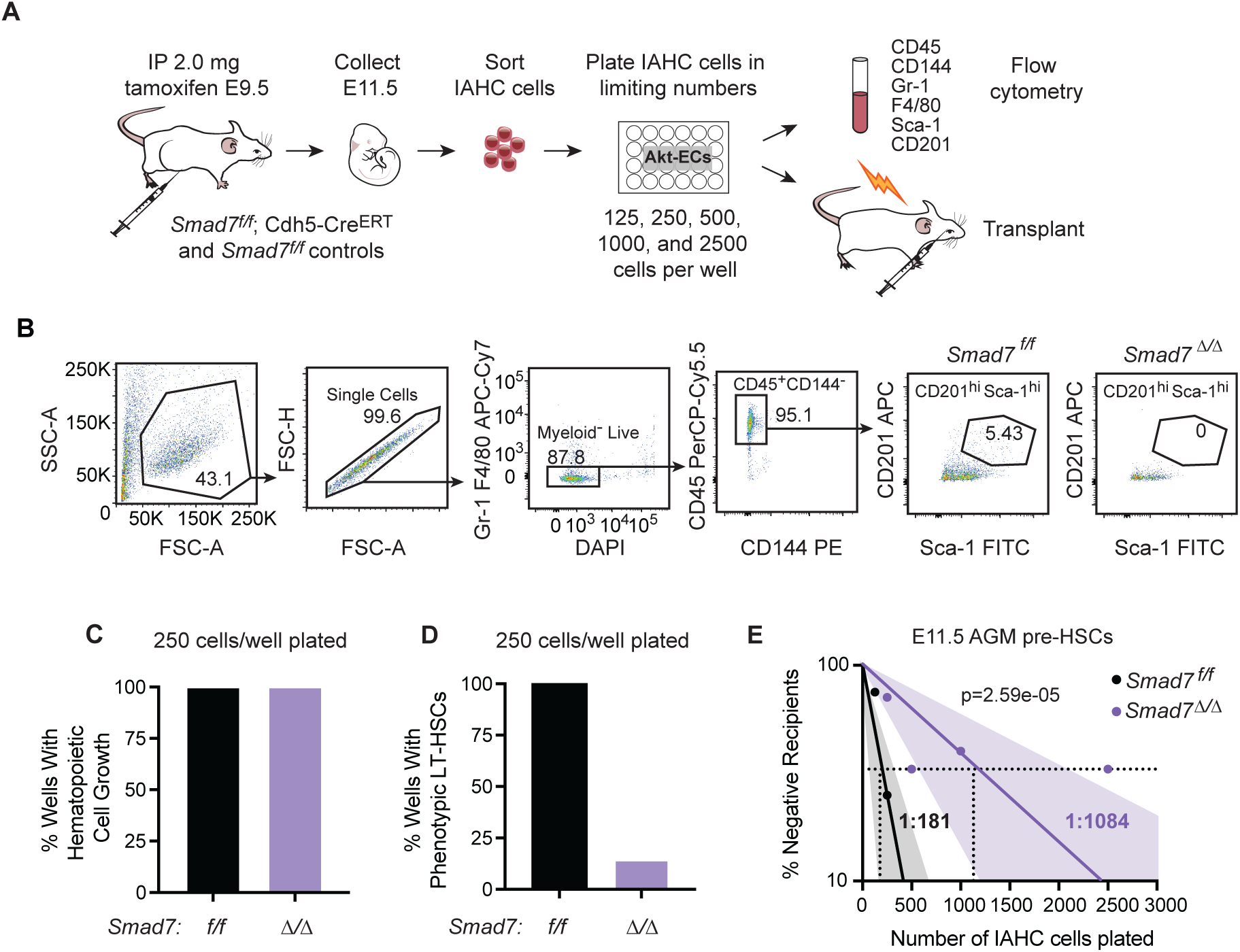
Loss of SMAD7 decreases the number of functional pre-HSCs. (A) Schematic of pre-HSC maturation assay. IAHC cells were purified from the AGM region of E11.5 *Smad7^f/f^* and *Smad7*^Δ*/*Δ^ embryos and plated in limiting numbers (125, 250, 500, 1000, and 2500 cells) on Akt-ECs and grown for 4 days. On day 4, half of the cells in each well were harvested for transplant and the other half was used for flow cytometry to identify phenotypic LT-HSCs. (B) Representative flow plots for the detection of phenotypic LT-HSCs in Akt-EC cultures. (C) Percentage of wells seeded with 250 IAHC cells that had evidence of hematopoietic cell growth defined as myeloid (Gr-1^+^ or F4/80^+^) or CD45^+^ cells by flow cytometry. (D) Percentage of wells seeded with 250 IAHC cells containing phenotypic LT-HSCs, defined as shown in panel B. (E) Quantification of LT-HSCs in pre-HSC maturation cultures determined by limiting dilution transplantation. Cells from each well were transplanted into CD45.1^+^ adult recipients (n=20-29 recipients with 3-8 recipients per dose for each genotype). The data represent 5 independent experiments. LT-HSC frequencies were calculated by ELDA^80^.

### SMAD7 is required for pre-HSC to HSC maturation

SMAD7 loss could impede the generation of pre-HSCs by decreasing the number of HECs fated to give rise to pre-HSCs or by impairing the ability of HECs to differentiate into pre-HSCs. Either of these scenarios would decrease the number of phenotypic pre-HSCs in *Smad7*^Δ*/*Δ^ embryos. To determine if pre-HSCs were efficiently generated, we isolated cells expressing endothelial markers and lacking differentiated blood cell markers (Ter119^-^CD41^lo/-^ CD31^+^CD144^+^ESAM^+^) from the caudal regions of E11.5 *Smad7*^Δ*/*Δ^ and control embryos (Fig. S4A) and performed single-cell RNA sequencing (scRNA-seq) (Fig. 4A). Summary statistics for collected cell populations are in Supplementary Table 1. We assigned cell identities using our previously published scRNA-seq data generated from multiple purified populations of cells collected from E9.5-E11.5 embryos (Fig. S4B) and projected data from the E11.5 *Smad7*^Δ*/*Δ^ and control samples onto a uniform manifold approximation and projection (UMAP) plot from the previously published dataset^10^ (Fig. 4A). Since the sorted E11.5 *Smad7*^Δ*/*Δ^ and control samples represent only a small fraction of the cell populations used to generate the original UMAP^10^, they occupy a limited portion of the UMAP (Fig. 4A). The relative proportion of IAHC cells, which includes pre-HSCs and hematopoietic cells that do not meet the threshold score for pre-HSCs (pre-HSC+other) relative to non-hemogenic endothelial cells (EC), was 25% lower in E11.5 *Smad7*^Δ*/*Δ^ embryos compared to control embryos (Fig. 4B). However, within the IAHC cell populations, the proportion of pre-HSCs relative to IAHC (other) cells was increased approximately 2-fold (33.3% in *Smad7*^Δ*/*Δ^ embryos versus 14.4% in *Smad7^f/f^* embryos) (Fig. 4C). The scRNA-seq data indicate that deletion of SMAD7 in endothelial cells does not reduce the number of molecularly defined pre-HSCs and instead increases their representation within the total population of IAHC cells.

**Figure 4.**
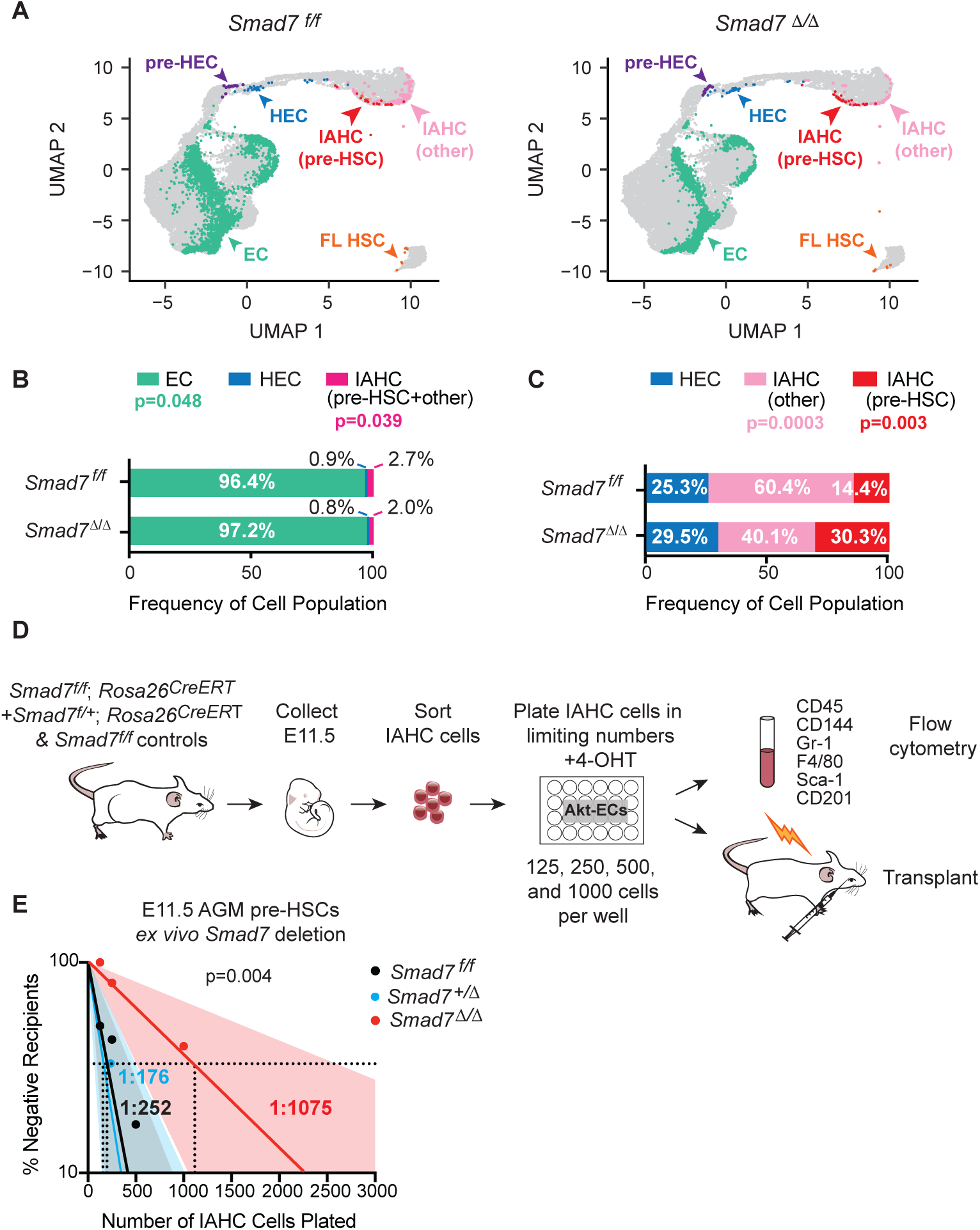
SMAD7 promotes the maturation of pre-HSCs into HSCs. (A) Uniform manifold approximation and projection (UMAP) plot of sorted CD31^+^CD44^+^CD41^low/-^c-Kit^low/+^ cells from E11.5 *Smad7^f/f^* and *Smad7*^Δ*/*Δ^ embryos projected onto the EHT trajectory from Zhu *et al*.^10^. Populations shown include pre-HEC, (immediate cell precursor of HEC); HEC; EC (endothelial cells excluding HEC and pre-HEC); IAHC (pre-HSC) (pre-HSCs within the IAHC population); IAHC (other) (IAHC cells that are not pre-HSCs); FL-HSC (E14.5 FL LSK CD48^-^CD150^+^ cells). (B) Distribution of cells between EC, HEC, and IAHC populations (including both pre-HSCs and other IAHC cells) in E11.5 *Smad7^f/f^*and *Smad7*^Δ*/*Δ^ embryos. *P* values indicate significant differences in the distribution of cells in specific populations in *Smad7^f/f^*and *Smad7*^Δ*/*Δ^ samples determined with the proportion test in R. (C) Distribution of cells with hematopoietic identity [HEC, IAHC (other), and IAHC (pre-HSC)] in E11.5 *Smad7^f/f^* and *Smad7*^Δ*/*Δ^ embryos. (D) Schematic for deleting *Smad7 ex vivo* in pre-HSCs. IAHC cells from E11.5 *Smad7 ^f/f^*, *Smad7 ^f/f^;Rosa26^CreERT+^*, and *Smad7^f/+^;Rosa26^CreERT+^* embryos were isolated and 4-OHT added when cells were plated on Akt-ECs. (E) Limiting dilution transplant from assay illustrated in panel D. n=15-28 recipients with 3-10 recipients per dose for each genotype. Data are from 3 independent experiments. HSC frequency calculated by ELDA^80^.

To directly examine whether SMAD7 loss specifically affected pre-HSC to HSC maturation, we allowed pre-HSCs to form normally, then deleted SMAD7. We used a ubiquitously expressed, tamoxifen-regulated Cre expressed from the *Rosa26* locus^49^ to delete SMAD7 during the pre-HSC to HSC maturation step. We generated E11.5 *Smad7^f/f^*; *Rosa26^CreERT^* embryos, purified IAHC cells (containing pre-HSCs) that were generated in the presence of SMAD7, plated them on Akt-ECs, then deleted SMAD7 *ex vivo* by adding 4-hydroxytamoxifen (4-OHT) to the cultures (Fig. 4D). *Ex vivo* deletion of SMAD7 in pre-HSCs resulted in an 87% decrease in functional LT-HSCs (Fig. 4E). Importantly, there was no decrease in the frequency of pre-HSCs in *Smad7^+/f^*; *Rosa26^CreERT^* control IAHC cells (*Smad7*^+*/*Δ^) able to mature into LT-HSCs, ruling out a non-specific effect of Cre^ERT^ activation or 4-OHT on pre-HSC to HSC maturation *ex vivo* (Fig. 4E). Therefore, SMAD7 is required for efficient pre-HSC to HSC maturation.

We analyzed the scRNA-seq data to gain insight into the mechanism underlying the pre-HSC to HSC maturation defect. Deletion of *Smad7* in endothelial cells caused large changes in gene expression in ECs, pre-HSCs, and IAHC (other) cells (there were too few HECs at E11.5 to analyze) (Fig. 5A,B). IAHC (other) cells from *Smad7*^Δ*/*Δ^ embryos, which would include LMPs, other progenitors, and hematopoietic cells at various stages of differentiation had the largest number of total genes up- and down-regulated compared to control embryos, while pre-HSCs had the smallest number of DEGs (Fig. 5A). Analysis of the genes up- and downregulated in ECs, IAHC (other) cells, and pre-HSCs in *Smad7*^Δ*/*Δ^ embryos showed very little overlap (Fig. 5B), indicating inhibition of TGFβ signaling results in unique transcriptional changes in different populations. Several of the top DEGs upregulated in *Smad7*^Δ*/*Δ^ pre-HSCs are involved in signaling cascades, cell cycle, and proliferation, such as *Kdr, Nr2f2,* and the cell cycle inhibitor *Cdkn1a* (Fig. 5C), while genes downregulated in *Smad7*^Δ*/*Δ^ pre-HSCs included those encoding transcription and translation factors, including known and predicted ribosomal subunits, such as *Rpl21, Rps27, Rpl9-ps6, Gm6133,* and the RNA polymerase I subunit *Taf1c* (Fig 5C). However, when we calculated a ribosome biogenesis score using all 322 genes belonging to the ribosome biogenesis pathway, there was no significant difference between control and *Smad7*^Δ*/*Δ^ pre-HSCs (Fig. 5D). Reactome analysis of *Smad7*^Δ*/*Δ^ pre-HSCs revealed enrichment for a broad variety of cell signaling pathways, including tyrosine kinase signaling, Notch4 activation, VEGFR signaling, and pathways related to the extracellular matrix, whereas genes downregulated in *Smad7*^Δ*/*Δ^ pre-HSCs were enriched for pathways related to genomic integrity (Fig. 5E). Genes upregulated in *Smad7*^Δ*/*Δ^ ECs were primarily related to translation, and terms for *Smad7*^Δ*/*Δ^ IAHC (other) cells included immune-related terms and signaling pathways (Fig. S5B). Thus, the divergent pathways affected by SMAD7 loss largely reflect the unique gene expression changes in ECs, pre-HSCs, and other IAHC cells.

**Figure 5.**
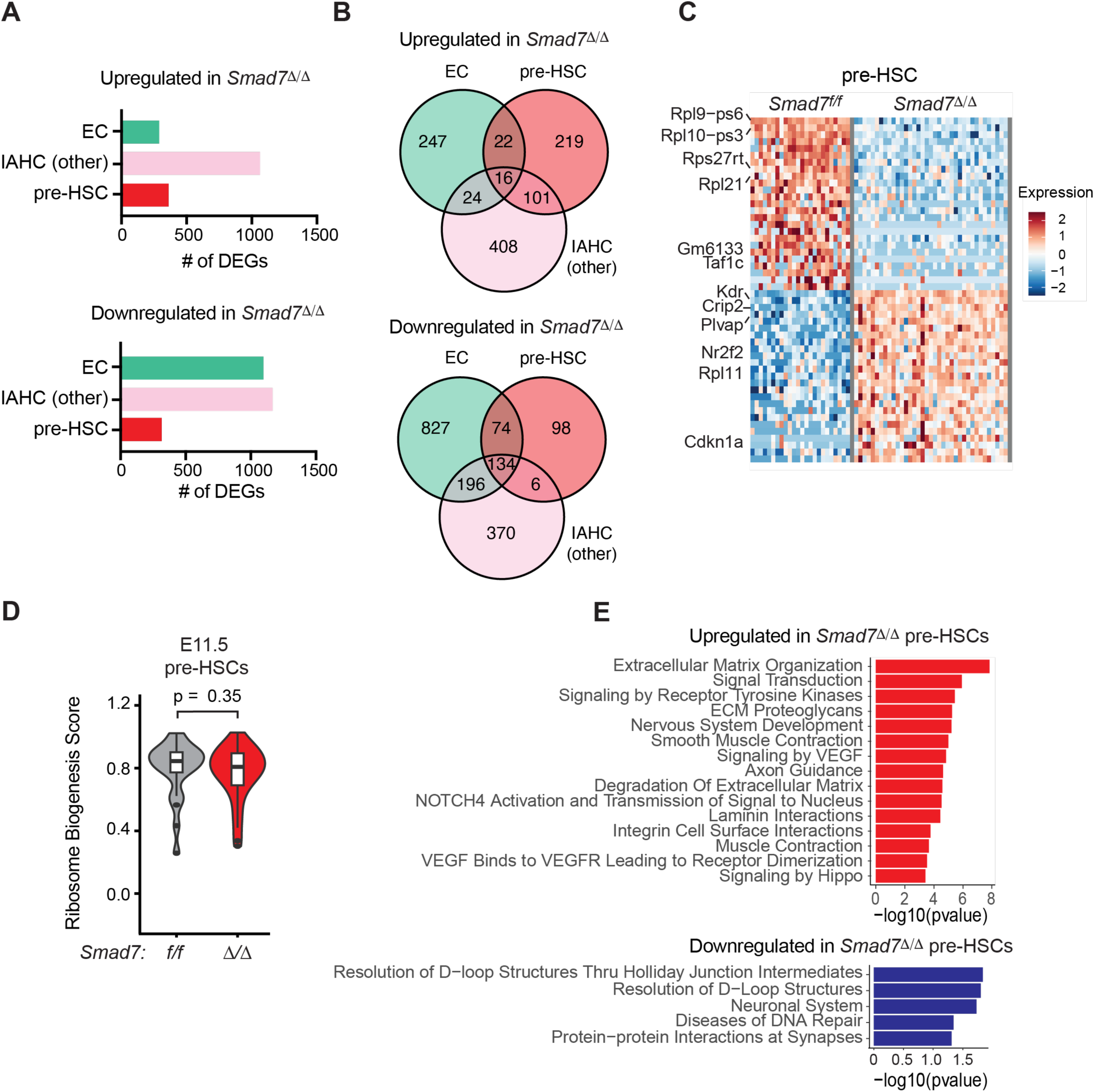
Loss of SMAD7 causes cell type-specific alterations in gene expression. (A) The total number of differentially expressed genes (DEGs) up- and down-regulated in E11.5 endothelial cells (ECs), IAHC (other) cells, and pre-HSCs. (B) Venn diagrams showing overlap of up- and down-regulated genes in *Smad7*^Δ*/*Δ^ cells for EC, IAHC (other), and pre-HSC populations. (C) Top 25 DEGs up- and down-regulated in *Smad7*^Δ*/*Δ^ pre-HSCs. (D) Ribosome biogenesis score for *Smad7^f/f^*and *Smad7*^Δ*/*Δ^ pre-HSCs using the average expression level of the ribosome biogenesis program and calculating a score using AddModuleScore in Seurat. (E) Pathways enriched in genes up- and down-regulated in *Smad7*^Δ*/*Δ^pre-HSCs determined by Reactome analysis.

### Loss of SMAD7 confers a selective advantage to fetal liver HSCs

A remaining possibility for why SMAD7 loss results in a severe reduction in embryonic LT-HSCs is that they could have a competitive disadvantage in transplant recipient mice. However, we found the opposite was true, as SMAD7-deficient LT-HSCs outcompeted LT-HSCs with intact or incompletely deleted *Smad7* alleles. For example, despite the 85% reduction in functional HSCs in the E12.5 A+U+V (Fig. 1I), the frequency of phenotypic LT-HSCs (LSK CD48^-^ CD150^+^) at E14.5 was similar in *Smad7*^Δ*/*Δ^ and control FLs (Fig. 6A), and the number of functional LT-HSCs was also equivalent (Fig. 6B). To confirm that E14.5 FL LT-HSCs and donor-derived LT-HSCs in transplant recipient mice had undergone biallelic *Smad7* deletion, we sorted LT-HSCs from the E14.5 FL and transplant recipient mice, plated them in methylcellulose, and genotyped individual colonies by PCR. The majority of colonies (53%) from E14.5 *Smad7*^Δ*/*Δ^ LT-HSCs (Lin^-^Sca-1^+^Kit^+^CD48^-^CD150^+^) cells contained two deleted *Smad7* alleles (*del/del*), 42% contained one deleted Smad7 allele (*f/del*), and 5% contained two intact *Smad7* floxed alleles (*f/f*) (Fig. 6C). In contrast, 89% of methylcellulose colonies from donor-derived LT-HSCs from transplant recipient mice contained two deleted *Smad7* alleles, demonstrating that *Smad7*^Δ*/*Δ^ LT-HSCs preferentially expanded in the recipients’ BM (Fig. 6C). The frequency of LT-HSCs with bi-allelic deletion of *Smad7* also increases in the fetal liver between E14.5 and E18.5 (53% at E14.5 vs 77% at E18.5), similar to the expansion of *Smad7*^Δ*/*Δ^ LT-HSCs following transplant (Fig. 6C). These data confirm that the defect in LT-HSC numbers occurs at the pre-HSC to HSC maturation step and is not caused by defective *in vivo* repopulating activity of *Smad7*^Δ*/*Δ^ LT-HSCs. On the contrary, it suggests that once *Smad7*^Δ*/*Δ^ LT-HSCs mature, they enjoy a selective advantage over LT-HSCs that escaped *Smad7* deletion.

**Figure 6.**
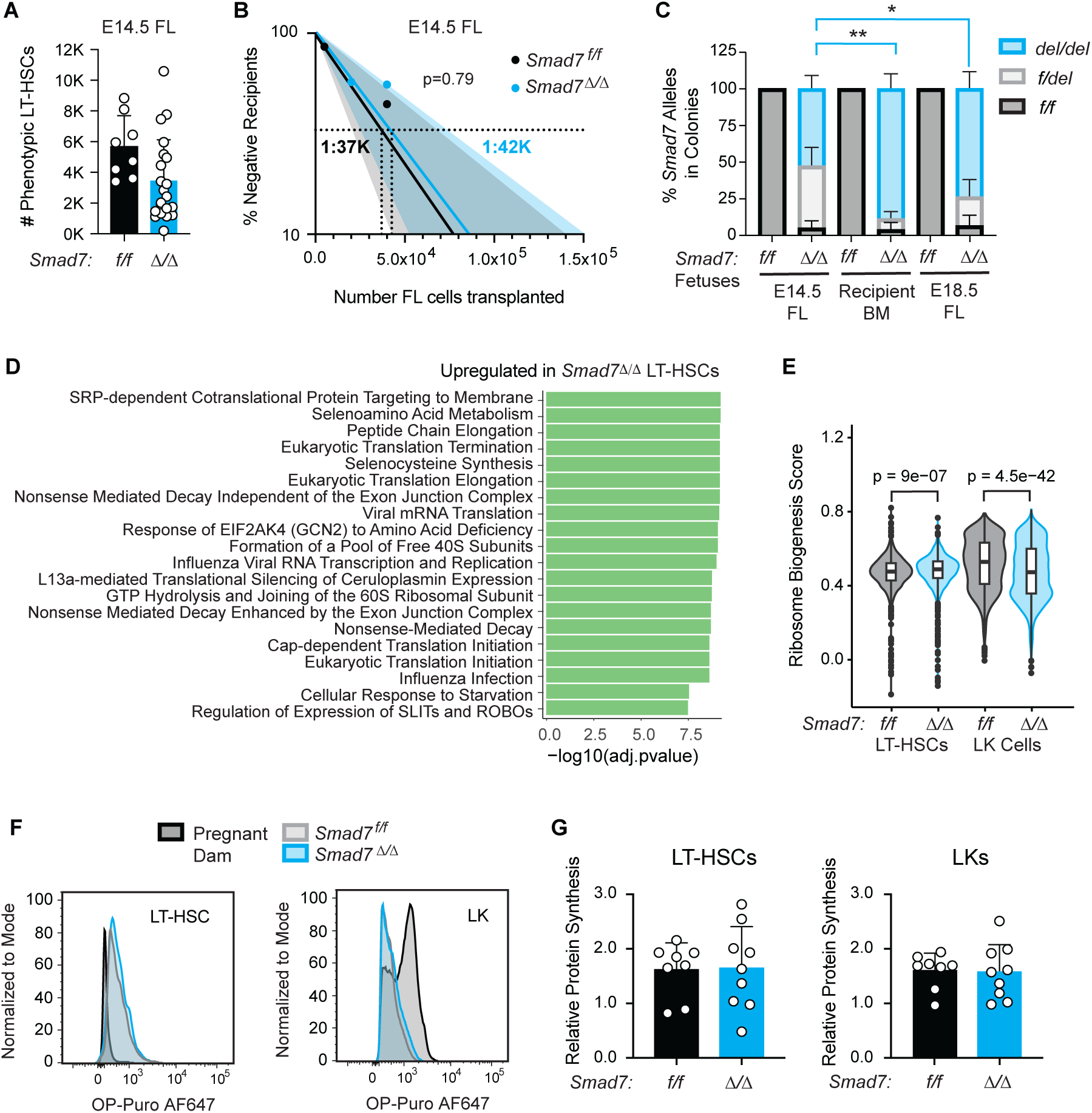
*Smad7*^Δ*/*Δ^ LT-HSCs preferentially expand in the fetal liver and bone marrow. (A) Quantification of phenotypic LT-HSCs (LSK CD48^-^ CD150^+^) in FLs of E14.5 *Smad7 ^f/f^* or *Smad7*^Δ*/*Δ^ fetuses by flow cytometry. Mean±SD, unpaired, two-tailed Student’s t-test, n=8-21 FLs and 3 independent experiments. (B) Similar frequencies of functional LT-HSCs in *Smad7^f/f^*and *Smad7*^Δ*/*Δ^ FLs as determined by limiting dilution analysis. 5K, 20K, 40K, 120K, or 400K unsorted *Smad7^f/f^* and *Smad7*^Δ*/*Δ^ FL cells (CD45.2^+^) were transplanted into lethally irradiated recipients (CD45.1^+^). Shown are percent negative recipients at each dose where negative was defined as <1% donor chimerism in the PB at 16 weeks. n= 38-39 recipients per genotype with 5-15 recipients per dose. (C) Percentage of colonies with biallelic (*del/del*), monoallelic (*f/del*) or no *Smad7* deletion (*f/f*) in E14.5 and E18.5 *Smad7^f/f^* and *Smad7*^Δ*/*Δ^ FLs, and in donor-derived LT-HSCs in the BM 16 weeks post-transplant of E14.5 FL cells. LT-HSCs were sorted from the FL or BM, plated in methylcellulose, and individual colonies were genotyped by PCR. Mean±SD, two-way ANOVA and Tukey’s multiple comparison test. n=24-153 individual colonies. (D) Pathways enriched in transcripts upregulated in E18.5 *Smad7*^Δ*/*Δ^ LT-HSCs using Reactome analysis. (E) Ribosome biogenesis scores in *Smad7 ^f/f^* or *Smad7*^Δ*/*Δ^ LT-HSCs and LK cells were calculated based on the average expression of 322 ribosome biogenesis pathway genes using AddModuleScore in Seurat. Wilcoxon Rank Sum Test. (F) Representative histograms from OP-Puro assay of phenotypic LT-HSCs and LK cells from E18.5 FLs. LT-HSCs from the pregnant dam were used as controls. (G) Relative protein synthesis rates measured by OP-Puro incorporation. One representative experiment is shown. Mean±SD, unpaired, two-tailed Student’s t-test, n=12-15 FLs and 3 independent experiments.

To investigate the mechanism driving the expansion of *Smad7*^Δ*/*Δ^ LT-HSCs, we performed scRNA-seq of E18.5 LT-HSCs and LK cells. We chose this time since by E18.5, 77% of LT-HSCs have biallelic Smad7 deletion (Fig. 6C), and tamoxifen injection of pregnant mice prevents successful delivery of live animals, making post-natal analysis more difficult. Reactome analysis of E18.5 *Smad7*^Δ*/*Δ^ LT-HSCs showed an enrichment in terms related to translation and a decrease in terms related to genomic integrity such as DNA repair, HDR, and cell cycle checkpoints (Fig. 6D, SFig. 6A). *Smad7*^Δ*/*Δ^ LT-HSCs have significantly higher ribosome biogenesis scores, while *Smad7*^Δ*/*Δ^ LK cells have lower ribosome biogenesis scores compared to their control counterparts (Fig. 6E). However, we could detect no difference in the rate of protein translation as measured by incorporation of O-propargyl-puromycin (OP-Puro) (Fig. 6F,G, SFig. 6B). Nevertheless, the GO analysis suggests that subtly increased translation could be one mechanism underlying the preferential expansion of *Smad7*^Δ*/*Δ^ LT-HSCs in the FL of E18.5 fetuses.

## Discussion

The maturation of pre-HSCs to HSCs is a poorly understood process in HSC formation that has been difficult to experimentally uncouple from the generation of pre-HSCs or LT-HSC function. Here, we definitively demonstrate that the TGFβ family signaling inhibitor SMAD7 is specifically required for pre-HSC to HSC maturation. We show that SMAD7 is not required for the generation of pre-HSCs, nor is it required for engraftment by FL LT-HSCs or their persistence in the FL or BM. SMAD7 is also not required for the generation of normal numbers of IAHCs or HSC-independent progenitors, including yolk sac-derived EMPs, LMPs, LMPPs, or TSPs. This is, to our knowledge the first gene shown to be necessary specifically for the efficient maturation of pre-HSCs into HSCs without broader requirements for hematopoietic ontogeny.

Multiple signaling pathways are active in HECs and involved in EHT^50^. scRNA-seq experiments suggest that a number of these pathways, including Notch, tumor necrosis factor, fluid shear stress, cytokine signaling, synthesis of eicosanoids, vitamins, sterols, Wnt, and TGFβ signaling are highly active in the immediate precursors of HECs (pre-HECs), and subsequently downregulated in HECs and IAHC cells^10,51,52^. In some cases, downregulation of a pathway was shown to impact HSC formation. For example, IAHC cells express high levels of the BMP4 inhibitor Noggin and lower levels of phosphorylated SMAD1/5/8 and SMAD2/3, the downstream effectors of BMP and TGFβ signaling, respectively, than aortic endothelial cells^37,53^. Addition of Noggin to AGM explant cultures improved their engraftment, indicating that downregulation of TGFβ family signaling promotes HSC formation^53^.

The contributions of downregulated EHT-related signaling to HSC formation have been most extensively detailed for the Notch pathway^54-60^. The receptors Notch1 and the delta-like ligand 4 (DLL4) are required for arterial specification^61-64^, a prerequisite for HSPC formation from arterial endothelium. DLL4 on IAHC cells activates Notch1 signaling in adjacent HECs and within IAHC cells themselves, with different outcomes. DLL4-mediated Notch1 signaling in HECs adjacent to the IAHCs enforces an arterial endothelial program that prevents HECs from undergoing EHT and joining the IAHCs^57,65^. This limits the number of HSCs^57^. Within IAHC cells, DLL4-mediated Notch1 signaling is thought to block cell cycle entry and hematopoietic differentiation^59^. Both activities of DLL4, on adjacent HECs and within the IAHC cells, are counteracted by JAGGED1 (JAG1)^59,65^. JAG1, which is a weaker stimulant of Notch1 activity than DLL4, downregulates endothelial genes and activates hematopoietic genes. JAG1 loss reduces the number of IAHC cells and HSCs by derepressing DLL4 signaling, enforcing arterial identity, and preventing HECs from joining the IAHCs^57,59,65,66^. Blocking JAG1 in IAHC cells upregulates genes involved in cell cycle entry and differentiation. JAG1 is thought to preserve an HSC-state in a subset of IAHC cells by cis-inhibiting DLL4-mediated signaling^59^.

The temporal requirement for downregulating TGFβ family signaling may be similar to the requirement for JAG1 cis-inhibition of DLL4-mediated Notch1 signaling in IAHCs, but not the earlier activity of JAG1 in promoting IAHC formation. Unlike JAG1 loss, SMAD7 loss does not affect the number of IAHC cells. Also, inappropriately elevated Notch signaling appears to affect the most immature populations of pre-HSCs, while our evidence suggests that SMAD7 loss affects more mature pre-HSCs. Pre-HSCs are categorized as less mature type I or more mature type II based on the absence or presence of cell surface CD45, respectively^24^. Inappropriately activating Notch signaling in pre-HSCs by culturing them on stromal cells ectopically expressing DLL1 will block the maturation of type I into type II pre-HSCs, but will not block the maturation of type II pre-HSCs into HSCs^58^. Although we did not explicitly address the role of *Smad7* in type I versus type II pre-HSCs, our data suggest that SMAD7 loss, at least in part, affects type II pre-HSCs. There are 4 times more type II than type I pre-HSCs at E11.5 in the AGM region^20^; therefore, an 85% decrease in pre-HSC to HSC maturation strongly suggests that loss of SMAD7 severely impairs the type II pre-HSC to HSC transition. We previously identified *Smad7* as being more highly expressed in E11.5 type II pre-HSCs (CD144^+^CD45^+^CD27^+^) versus LMPs^10^, consistent with a role for SMAD7 in type II pre-HSCs. Also, our current studies show an increased proportion of molecular type II pre-HSCs in the absence of SMAD7, further suggesting that inhibition of TGFβ family signaling is most critical for the transition from type II pre-HSCs to HSCs. Finally, it was previously reported that several BMP4 target genes are expressed at significantly lower levels in type II pre-HSCs than in type I pre-HSCs^53^. The TGFβ/BMP pathway inhibitor *Pdzk1ip1* ^67^ is also expressed in E11.5 type II pre-HSCs^10,30^, and *Smad6* is highly expressed in HSC-primed HECs and IAHC cells^37,59^. Thus, multiple inhibitors may participate in restraining TGFβ family signaling in the precursors of HSCs.

Once HSCs have matured from pre-HSCs, loss of SMAD7 confers a selective advantage in the FL and adult BM. We have not yet discerned the mechanism behind this switch in SMAD7’s function from promoting pre-HSC maturation to restraining LT-HSC expansion. Relatively little is known about SMAD7’s role in FL and BM LT-HSCs. Retroviral overexpression of SMAD7 in BM cells reduced proliferation and increased LT-HSC self-renewal^68^, opposite to what we would have predicted based on our results, but the methodologies used (overexpression versus conditional deletion) in our studies are very different. In our approach, SMAD7 was also deleted in endothelial cells, which are an essential component of LT-HSC niches^69,70^, although at least some aspect of the selective LT-HSC expansion is cell autonomous, as it also occurred in transplant recipient mice. The most prominent Reactome terms associated with genes upregulated in E18.5 *Smad7*^Δ*/*Δ^ LT-HSCs were related to protein translation, and the ribosome biogenesis score was significantly higher. We did not detect a difference in protein synthesis rates in *Smad7*^Δ*/*Δ^ LT-HSCs *in vivo*, although we cannot rule out a subtle change that is difficult to detect experimentally. TGFβ and BMP signaling regulate multiple HSC properties, including proliferation, quiescence, self-renewal, and apoptosis ^71-75^. Further experiments are necessary to determine why *Smad7*^Δ*/*Δ^ FL LT-HSCs selectively expand, whether through a proliferative advantage, a decrease in apoptosis, or skewing towards symmetric self-renewal divisions^76^.

The different effects of TGFβ/BMP signaling at distinct stages of hematopoietic ontogeny provide a rationale for further modulating their activity in protocols designed to optimize LT-HSC production from induced pluripotent stem cells (iPSCs). Indeed, a recent landmark study describing a protocol for producing robust multi-lineage engraftment of immunodeficient mice by human iPSC-derived HSCs uses precisely this approach^77^. The protocol utilizes BMP4 to specify mesoderm, then a TGFβ inhibitor to pattern AGM-like mesoderm expressing *HOXA* genes, then adds BMP4 again to promote endothelial cell formation and hematopoietic development^77^. Our data suggest that additional temporal modifications of TGFβ and/or BMP signaling, i.e., lowering one or both pathway activities to promote pre-HSC to HSC maturation, and later increasing them to promote HSC expansion may further improve the yield of LT-HSCs from iPSCs.

## Methods

### Mouse strains and embryos

*Smad7^f/f^*(B6.Cg-*Smad7^tm1.1Ink^*/J; 017008) and B6.129-*Gt(ROSA)26Sor^tm1(cre/ERT2)Tyj^*/J (008463) mice^49^ were purchased from Jackson Laboratory. *Cdh5-* Cre^ERT^ (Tg(Cdh5-cre/ERT2)1Rha) mice were a gift from Ralf Adams (Max Planck Institute for Molecular Biomedicine, Münster, Germany)^40^. All mice were backcrossed 6-10 generations to C57BL/6J. For embryo generation, the morning after mating is considered E0.5. For *in vivo* deletion of floxed alleles, tamoxifen (2mg/mouse in corn oil) was given to pregnant dams on E9.5 by intraperitoneal injection. E10.5-E11.5 embryos were staged on the day of harvest by counting somite pairs. The yolk sac was collected for genotyping of individual embryos. Embryos that were underdeveloped or abnormal were discarded. Mice were handled according to protocols approved by the University of Pennsylvania’s Institutional Animal Care and Use Committee.

### Confocal microscopy

Embryos were prepared for whole-mount microscopy as previously described^78^. A Zeiss LSM 880 AxioObserver inverted confocal microscope with ZEN 2.3 SP1 software was used to acquire Z-optical projections. Images were processed using Fiji software^79^. All primary antibodies were used at a concentration of 1:500 except for c-Kit and Cytokeratin 8, which were used at concentrations of 1:250 and 1:20, respectively. Secondary antibodies were all used at a concentration of 1:1000. A complete list of primary and secondary antibodies used can be found in Supplementary Table 2. To measure RUNX1 intensity in HECs and IAHC cells, we calculated Corrected Total Cell Fluorescence (CTCF) using Fiji software. CTCF was defined as [integrated density of the cell - (area of the selected cell × mean intensity of background)]. A power analysis indicated that to detect a 10% change in RUNX1 intensity with a significance level of P≤0.01, a sample size of 220 cells per genotype was required.

### Flow cytometry and cell sorting

All antibodies are in Supplementary Table 2 and were used at a dilution of 1:200. Either DAPI or LIVE/DEAD Fixable Aqua (ThermoFisher L34957) was used to exclude dead cells. Embryos were prepared for flow cytometry by removing the head, cardiac and pulmonary regions, liver, digestive tube, tail, and limb buds. The remaining tissue containing the aorta-gonad-mesonephros (AGM) region, parts of the body wall, somites, and umbilical and vitelline vessels was collected. Tissues were dissected in phosphate-buffered saline (PBS)/10% fetal bovine serum (FBS) and penicillin/streptomycin (Sigma). After dissociation in 0.125% collagenase (Sigma) for 1 hour, tissues were mechanically dissociated, washed, and filtered through a 100 μm filter to create a single-cell suspension. Cells were immediately stained with an antibody cocktail for 45-60 minutes at 4°C before being washed and resuspended in FACS buffer (PBS + 2% FBS). E12.5, E14.5, and E18.5 fetal livers were removed from the embryo, mechanically dissociated by pipetting, washed, and filtered through a 100 μm filter and stained as described. Embryonic cells were sorted on a BD Influx with a 100 μm nozzle at flow rates less than 4,000 events/second and collected directly into either PBS+20% FBS (single-cell RNA sequencing) or X-vivo 20 (IAHCs for pre-HSC maturation). E14.5 and E18.5 fetal liver cells and adult bone marrow were sorted on a BD FACSAria II with a 70 μm nozzle. Flow cytometry was performed on BD LSRII cytometers and data were analyzed using FlowJo software (v10.8.1, BD Biosciences).

### Limiting dilution LMP assay

OP9 stromal cells were cultured in αMEM supplemented with 20% FBS and antibiotics and plated into 96-well plates at a density of 5,500 cells/well one day before adding dilutions of dissociated embryonic tissues. The medium was supplemented with 5 ng/ml Flt3L and 10 ng/ml IL-7 (Peprotech). Cells were collected and analyzed by flow cytometry 7-10 days after culture. B cells were identified as CD19^+^ B220^+^ and myeloid cells as Gr-1^+^ CD11b^+^. Progenitor frequencies were calculated using extreme limiting dilution analysis (ELDA)^80^. OP9s were purchased from ATCC and were mycoplasma negative.

### Hematopoietic stem cell transplants

B6.SJL-*Ptprc^a^Pepc^b^*/BoyJ (CD45.1) mice were subjected to a split dose of lethal irradiation (900 cGy) 4 hours apart. For AGM transplants at E11.5 and E12.5, recipients received either 0.1, 0.3, or 1 embryo equivalent (ee) of a disassociated donor AGM region (CD45.2) with 2.5×10^5^ splenocytes (CD45.1/CD45.2) by retro-orbital injection. For FL transplants at E12.5, recipients received 0.1, 0.05, or 0.02 (1/10^th^, 1/20^th^, and 1/50^th,^ respectively) of donor FL cells (CD45.2) with 2.5 × 10^5^ splenocytes (CD45.1/CD45.2) by retro-orbital injection. For E14.5 transplants, each recipient received 5K, 20K, 40K, 120K, or 400K donor FL cells (CD45.2) with 2.5 × 10^5^ splenocytes (CD45.1/CD45.2) by retro-orbital injection. Donor (CD45.2) engraftment was assessed in PB at weeks 4, 8, 12, and 16, and in BM at week 16 post-transplantation. LT-HSC frequencies were determined by ELDA^80^.

### Pre-HSC ex vivo maturation assays

pre-HSC cultures were performed as previously described^47^. Briefly, endothelial cells constitutively expressing Akt (Akt-ECs) are maintained on plates coated with 0.1% gelatin in Akt-EC Media [IMDM supplemented with 20% FBS (Gibco)], penicillin/streptomycin, L-glutamine, heparin (0.1 mg/ml; H3149-100KU, Sigma-Aldrich), endothelial cell growth supplement (100 μg/ml; 02-101, Millipore Sigma). One day before plating primary embryonic cells, Akt-ECs are plated at a density of 30,000 Akt-ECs/well on gelatin-coated 24-well tissue culture plates in Akt-EC Media. Akt-EC Media was changed to serum-free X-vivo 20 media (BW04-448Q, Fisher Scientific) containing 100ng/mL SCF, IL-3, Flt-3, and 20ng/mL TPO (all cytokines from Peprotech) 8-12 hours before adding embryonic cells. Sorted IAHC cells are plated in limiting numbers (125, 250, 500, 1000, or 2500 cells/well) on Akt-ECs and cultured for 4 days. For all Rosa26^CreERT^ pre-HSC maturation assays, 4-OHT (1 μM; H6278, Sigma-Aldrich) was added to all wells when IAHC cells were plated. After 4 days, wells were treated with accutase for 5-10 minutes to lift IAHCs and Akt-ECs for analysis by flow cytometry and transplant into lethally irradiated mice (B6.SJL-*Ptprc^a^Pepc^b^*/BoyJ).

### Colony assays and single colony genotyping

Yolk sacs were harvested and digested with 0.125% collagenase for 1 hour at 37°C. 1/40th of each yolk sac was added to methylcellulose (Stem Cell Technologies, M3434) and plated. Colonies were scored after 7 days. For single colony genotyping, either one entire AGM region or 500 donor LT-HSCs were plated, and single colonies were picked and assayed by PCR for *Smad7^f^* and *Smad7*^Δ^ (deleted) alleles. AGMs were flushed with PBS to remove circulating cells before plating. Primers for *Smad7^f^* and *Smad7*^Δ^ (deleted) alleles were previously described^81^.

### Single Cell RNA Sequencing

We performed whole transcriptome analysis using the BD Rhapsody Single-Cell Analysis System according to the manufacturer’s instructions (BD Biosciences). In short, approximately 15,000 sorted cells were loaded onto a BD Rhapsody cartridge (BD Biosciences, 633733), and single cells were captured by gravity. The low cell density compared to the number of microwells in the cartridge and visual scanning of the cartridge after loading assured single-cell occupancy of wells. Cells were lysed in microwells, and mRNA molecules were allowed to hybridize to barcoded capture oligos on beads, which were then collected from the microwells for cDNA synthesis and library prep with the BD Whole Transcriptome Analysis Amplification Kit (BD Biosciences, 665915). Libraries were sequenced on an Illumina NovaSeq 6000. Fastq files were processed with the BD Rhapsody analysis pipeline (BD Biosciences) on Seven Bridges (https://www.sevenbridges.com) according to the manufacturer’s recommendations.

### Analysis of scRNA-seq from the BD Rhapsody Single-Cell system

The raw cell-by-gene count matrix was first normalized using Monocle 3^82^ and then converted into Seurat^83^ objects for *Smad7^f/f^* (WT) and *Smad7*^Δ*/*Δ^ (KO) samples, respectively. Subsequently, we selected the top 2,000 variably expressed genes (VEGs) using the *FindVariableFeatures* function in Seurat with the default setting. The VEGs were further scaled by the ScaleData function in Seurat. We then performed principal component analysis (PCA) using the *RunPCA* function in Seurat with 20 principal components (PCs). Uniform Manifold Approximation and Projection (UMAP) was calculated using the *RunUMAP* function in Seurat for visualization with the default setting. Cells in each dataset were projected onto the reference EHT trajectory established by Zhu *et al.*^10^, following a pipeline described in Chen *et al.*^84^. Specifically, for each of the VEGs in the reference Seurat object, we extracted mean and standard deviation of the normalized data. Then, for each corresponding gene in the query data, we scaled the data with the corresponding mean and standard deviation of the reference data. This transformed gene-by-cell matrix was then multiplied by the PCA feature loading matrix from the reference Seurat object to calculate the projected PC scores for each query cell. Finally, the umap_transfer function from the uwot R package was used to generate the projected UMAP coordinates for each query cell. The cell type of each query cell was defined as the cell type of the nearest (in Euclidean distance) reference cell in the UMAP embedding. For each of the pre-HSC, EC, and IAHC (other) populations, we performed differential gene expression analysis between *Smad7^f/f^* and *Smad7*^Δ*/*Δ^ cells using the *FindAllMarkers* function in Seurat, with the parameters test.use = "LR", logfc.threshold = 0.25, and max.cells.per.ident = 200. To visualize the top differentially expressed genes (DEGs), we generated heatmaps for each comparison using the *DoHeatmap* function in Seurat. Enriched REACTOME pathways for the DEGs were identified using the enrichR R package.

### Analysis of scRNA-seq from 10x genomics platform (LT-HSC)

The raw data were first preprocessed using cellranger (v8.0.0) with mouse genome assembly GRCm39, followed by additional processing using customized R scripts. Specifically, we filtered out cells with fewer than 1000 UMIs or greater than 50000 UMIs, as well as cells with fewer than 500 genes or greater than 10000 genes. Cells with more than 5% of UMI from mitochondrial genes were removed for downstream analyses. We then performed the standard Seurat pipeline as above for the scRNA-seq from the BD Rhapsody Single-cell system. In addition, we calculated the Ribosome biogenesis score for each cell using *AddModuleScore* function in Seurat with all 322 genes belong to ‘ribosome biogenesis’ pathway from https://www.informatics.jax.org/vocab/gene_ontology/GO:0042254.

### Statistics

All statistics were performed using Prism software (v. 9.0.0). An unpaired two-tailed Student’s *t*-test was used for pairwise comparisons. One-way ANOVA with Tukey’s multiple comparison test was used for >2 groups. For all *t*-tests and one-way ANOVAs, \**P*<0.05, \*\**P*<0.01, \*\*\**P*<0.001 and \*\*\*\**P*<0.0001. Fisher’s exact test was used for the categorical analysis of variables where \**P*<0.0332, \*\**P*<0.0021, \*\*\**P*<0.0002, \*\*\*\**P*<0.0001.

## Data Availability

All datasets will be made publicly available on GEO.

## Acknowledgments

We thank Andrea Stout and AJ Lucy at the Cell and Developmental Biology Microscopy Core for confocal microscopy assistance. We thank Jennifer Jakubowski, Andrew Morschauser, William Murphy, Charles Pletcher, and Shifu Tian at the Penn Cytometrics and Cell Sorting Resource Laboratory for their assistance. We thank Michael Bowman for assistance with initial computational analyses.

## Author contributions

N.A.S. and L.F.B conceived the project, designed experiments, and wrote the manuscript.

L.F.B, H.H.A, and J.T. carried out experiments and analyzed the data. W.Y. performed computational analyses. C.H.C. performed scRNA-seq experiments. K.T. provided guidance on the project. N.A.S and K.T. acquired funding for the project.

## Figure Legends

**Supplementary Figure 1. Loss of SMAD7 at E9.5 does not affect EMP numbers.** (A) Quantification of EMPs per yolk sac in *Smad7 ^f/f^* or *Smad7*^Δ*/*Δ^ embryos at E10.5. Colonies were scored on day 7 after plating in methylcellulose. Mean±SD. Two-way ANOVA and Šidák’s test for multiple comparisons. GEMM, granulocyte, erythrocyte, monocyte, megakaryocyte progenitors; BFU-E, burst-forming unit-erythroid progenitors; GM, granulocyte-monocyte progenitors. (B) Individual colonies from EMPs grown in methylcellulose were picked, and deletion of the *Smad7* alleles was assessed by PCR. Embryo genotypes are shown on the bottom and colony genotypes (*del/del*, biallelic *Smad7* deletion; *f/del*, monoallelic *Smad7* deletion; *f/f*, no *Smad7* deletion) on the right. (C) Representative confocal images of HECs (indicated by asterisks) and IAHC cells (indicated by white arrowheads) in the dorsal aorta of E10.5 embryos stained for CD31 (red) and RUNX (green). Scale bar=50µM. Related to Figure 1.

**Supplementary Figure 2. *Smad7* deletion decreases the number of multi-lineage repopulating LT-HSCs in the AGM region.** (A) Gating strategy for analyzing donor contribution to the PB of primary transplant recipient mice. Mice were engrafted with E12.5 A+U+V cells. Shown are representative examples of recipient mice with multilineage engraftment, and lymphoid-only engraftment in PB. (B) Gating strategy for identifying phenotypic LT-HSCs in the BM of transplant recipient mice. (C) Contribution of donor-derived E12.5 A+U+V cells from *Smad7*^Δ^*^/^*^Δ^ embryos to phenotypic CD45.2^+^ LT-HSCs in the BM of primary transplant recipient mice. (D) Contribution of donor-derived E12.5 A+U+V cells from *Smad7*^Δ^*^/^*^Δ^ embryos to phenotypic Lin^-^ cells in the BM of primary transplant recipient mice. Related to Figure 1.

**Supplementary Figure 3. Gating strategy for isolating IAHC cells.** Representative flow plots showing gating strategy for isolating E11.5 IAHC cells used in pre-HSC limiting dilution assays. Related to Figure 3.

**Supplementary Figure 4. SMAD7 does not influence pre-HSC formation.** (A) Representative flow plots showing the gating strategy for sorting ECs, HECs, and IAHC cells from E11.5 *Smad7^f/f^*and *Smad7*^Δ^*^/^*^Δ^ embryos (*Smad7*^Δ^*^/^*^Δ^ = *Smad7^f/f^*; Cdh5-Cre^ERT^) for scRNA-seq. 7-8 embryos were pooled for each genotype. (B) Uniform manifold approximation and projection (UMAP) plot of scRNA-seq data from sorted cells of E11.5 *Smad7^f/f^* and *Smad7*^Δ^*^/^*^Δ^ embryos. Cell types (left) were assigned by projecting the data onto the EHT trajectory from Zhu *et al.*^10^. Each cell was then assigned the annotation of its nearest neighbor in the reference trajectory. The IAHS (middle) and pre-HSC (right) scores were calculated using the *AddModuleScore* function in Seurat. This calculation was based on the top 100 upregulated genes for each respective population, downloaded from Zhu *et al.*^10^. Related to Figure 4.

**Supplementary Figure 5. Loss of SMAD7 causes broad dysregulation of gene programs in all populations.** (A) Top 25 differentially expressed genes (DEGs) up- or downregulated in *Smad7*^Δ*/*Δ^ EC and IAHC (other) cells. (B) Pathways up- or downregulated in ECs and IAHC (other) cells in *Smad7*^Δ*/*Δ^ embryos by Reactome analysis. Related to Figure 5.

**Supplementary Figure 6. Loss of SMAD7 causes downregulation of DNA repair genes in E18.5 *Smad7***^Δ***/***Δ^ **LT-HSCs.** Reactome pathway enrichment showing downregulated pathways in *Smad7*^Δ*/*Δ^ LT-HSCs at E18.5. Related to Figure 6.

## References

1. Soares-da-Silva, F., Elsaid, R., Peixoto, M.M., Nogueira, G., Pereira, P., Bandeira, A., and Cumano, A. (2023). Assembling the layers of the hematopoietic system: A window of opportunity for thymopoiesis in the embryo. Immunol Rev 315, 54–70. 10.1111/imr.13187.

2. Kissa, K., and Herbomel, P. (2010). Blood stem cells emerge from aortic endothelium by a novel type of cell transition. Nature 464, 112–115. nature08761 [pii] 10.1038/nature08761.

3. Hadland, B., and Yoshimoto, M. (2018). Many layers of embryonic hematopoiesis: new insights into B-cell ontogeny and the origin of hematopoietic stem cells. Exp Hematol 60, 1–9. 10.1016/j.exphem.2017.12.008.

4. Palis, J., Robertson, S., Kennedy, M., Wall, C., and Keller, G. (1999). Development of erythroid and myeloid progenitors in the yolk sac and embryo proper of the mouse. Development 126, 5073–5084.

5. Frame, J.M., Fegan, K.H., Conway, S.J., McGrath, K.E., and Palis, J. (2015). Definitive Hematopoiesis in the Yolk Sac Emerges from Wnt-Responsive Hemogenic Endothelium Independently of Circulation and Arterial Identity. Stem cells. 10.1002/stem.2213.

6. Godin, I., Dieterlen-Lièvre, F., and Cumano, A. (1995). Emergence of multipotent hematopoietic cells in the yolk sac and paraaortic splanchnopleura of 8.5 dpc mouse embryos. Proc. Natl. Acad. Sci. USA 92, 773-777.

7. Yokota, T., Huang, J., Tavian, M., Nagai, Y., Hirose, J., Zuniga-Pflucker, J.C., Peault, B., and Kincade, P.W. (2006). Tracing the first waves of lymphopoiesis in mice. Development 133, 2041–2051. dev.02349 [pii] 10.1242/dev.02349.

8. Li, Y., Esain, V., Teng, L., Xu, J., Kwan, W., Frost, I.M., Yzaguirre, A.D., Cai, X., Cortes, M., Maijenburg, M.W., et al. (2014). Inflammatory signaling regulates embryonic hematopoietic stem and progenitor cell production. Genes Dev 28, 2597–2612. 10.1101/gad.253302.114.

9. Cumano, A., Dieterlen-Lièvre, F., and Godin, I. (1996). Lymphoid potential, probed before circulation in mouse, is restricted to caudal intraembryonic splanchnopleura. Cell 86, 907–916.

10. Zhu, Q., Gao, P., Tober, J., Bennett, L., Chen, C., Uzun, Y., Li, Y., Howell, E.D., Mumau, M., Yu, W., et al. (2020). Developmental trajectory of prehematopoietic stem cell formation from endothelium. Blood 136, 845–856. 10.1182/blood.2020004801.

11. Ohmura, K., Kawamoto, H., Fujimoto, S., Ozaki, S., Nakao, K., and Katsura, Y. (1999). Emergence of T, B, and myeloid lineage-committed as well as multipotent hemopoietic progenitors in the aorta-gonad-mesonephros region of day 10 fetuses of the mouse. J Immunol 163, 4788-4795. ji_v163n9p4788 [pii].

12. Boiers, C., Carrelha, J., Lutteropp, M., Luc, S., Green, J.C., Azzoni, E., Woll, P.S., Mead, A.J., Hultquist, A., Swiers, G., et al. (2013). Lymphomyeloid contribution of an immune-restricted progenitor emerging prior to definitive hematopoietic stem cells. Cell Stem Cell 13, 535–548. 10.1016/j.stem.2013.08.012.

13. Luis, T.C., Luc, S., Mizukami, T., Boukarabila, H., Thongjuea, S., Woll, P.S., Azzoni, E., Giustacchini, A., Lutteropp, M., Bouriez-Jones, T., et al. (2016). Initial seeding of the embryonic thymus by immune-restricted lympho-myeloid progenitors. Nat Immunol 17, 1424–1435. 10.1038/ni.3576.

14. Owen, J.J., and Ritter, M.A. (1969). Tissue interaction in the development of thymus lymphocytes. J Exp Med 129, 431–442. 10.1084/jem.129.2.431.

15. Elsaid, R., Meunier, S., Burlen-Defranoux, O., Soares-da-Silva, F., Perchet, T., Iturri, L., Freyer, L., Vieira, P., Pereira, P., Golub, R., et al. (2021). A wave of bipotent T/ILC-restricted progenitors shapes the embryonic thymus microenvironment in a time-dependent manner. Blood 137, 1024–1036. 10.1182/blood.2020006779.

16. Ramond, C., Berthault, C., Burlen-Defranoux, O., de Sousa, A.P., Guy-Grand, D., Vieira, P., Pereira, P., and Cumano, A. (2014). Two waves of distinct hematopoietic progenitor cells colonize the fetal thymus. Nat Immunol 15, 27–35. 10.1038/ni.2782.

17. Dignum, T., Varnum-Finney, B., Srivatsan, S.R., Dozono, S., Waltner, O., Heck, A.M., Ishida, T., Nourigat-McKay, C., Jackson, D.L., Rafii, S., et al. (2021). Multipotent progenitors and hematopoietic stem cells arise independently from hemogenic endothelium in the mouse embryo. Cell reports 36, 109675. 10.1016/j.celrep.2021.109675.

18. Inlay, M.A., Serwold, T., Mosley, A., Fathman, J.W., Dimov, I.K., Seita, J., and Weissman, I.L. (2014). Identification of multipotent progenitors that emerge prior to hematopoietic stem cells in embryonic development. Stem cell reports 2, 457–472. 10.1016/j.stemcr.2014.02.001.

19. Patel, S.H., Christodoulou, C., Weinreb, C., Yu, Q., da Rocha, E.L., Pepe-Mooney, B.J., Bowling, S., Li, L., Osorio, F.G., Daley, G.Q., and Camargo, F.D. (2022). Lifelong multilineage contribution by embryonic-born blood progenitors. Nature 606, 747–753. 10.1038/s41586-022-04804-z.

20. Rybtsov, S., Ivanovs, A., Zhao, S., and Medvinsky, A. (2016). Concealed expansion of immature precursors underpins acute burst of adult HSC activity in foetal liver. Development 143, 1284–1289. 10.1242/dev.131193.

21. Müller, A.M., Medvinsky, A., Strouboulis, J., Grosveld, F., and Dzierzak, E. (1994). Development of hematopoietic stem cell activity in the mouse embryo. Immunity 1, 291–301.

22. Kumaravelu, P., Hook, L., Morrison, A.M., Ure, J., Zhao, S., Zuyev, S., Ansell, J., and Medvinsky, A. (2002). Quantitative developmental anatomy of definitive haematopoietic stem cells/long-term repopulating units (HSC/RUs): role of the aorta-gonad-mesonephros (AGM) region and the yolk sac in colonisation of the mouse embryonic liver. Development 129, 4891–4899.

23. Medvinsky, A., and Dzierzak, E. (1996). Definitive hematopoiesis is autonomously initiated by the AGM region. Cell 86, 897–906.

24. Rybtsov, S., Sobiesiak, M., Taoudi, S., Souilhol, C., Senserrich, J., Liakhovitskaia, A., Ivanovs, A., Frampton, J., Zhao, S., and Medvinsky, A. (2011). Hierarchical organization and early hematopoietic specification of the developing HSC lineage in the AGM region. J Exp Med 208, 1305–1315. jem.20102419 [pii] 10.1084/jem.20102419.

25. Taoudi, S., Gonneau, C., Moore, K., Sheridan, J.M., Blackburn, C.C., Taylor, E., and Medvinsky, A. (2008). Extensive hematopoietic stem cell generation in the AGM region via maturation of VE-cadherin+CD45+ pre-definitive HSCs. Cell Stem Cell 3, 99–108. S1934-5909(08)00283-X [pii] 10.1016/j.stem.2008.06.004.

26. Bennett, L.F., Mumau, M.D., Li, Y., and Speck, N.A. (2022). MyD88-dependent TLR signaling oppositely regulates hematopoietic progenitor and stem cell formation in the embryo. Development 149. 10.1242/dev.200025.

27. Yokomizo, T., Ideue, T., Morino-Koga, S., Tham, C.Y., Sato, T., Takeda, N., Kubota, Y., Kurokawa, M., Komatsu, N., Ogawa, M., et al. (2022). Independent origins of fetal liver haematopoietic stem and progenitor cells. Nature 609, 779–784. 10.1038/s41586-022-05203-0.

28. Kieusseian, A., Brunet de la Grange, P., Burlen-Defranoux, O., Godin, I., and Cumano, A. (2012). Immature hematopoietic stem cells undergo maturation in the fetal liver. Development 139, 3521–3530. dev.079210 [pii] 10.1242/dev.079210.

29. Ema, H., and Nakauchi, H. (2000). Expansion of hematopoietic stem cells in the developing liver of a mouse embryo. Blood 95, 2284–2288.

30. Hadland, B., Varnum-Finney, B., Dozono, S., Dignum, T., Nourigat-McKay, C., Heck, A.M., Ishida, T., Jackson, D.L., Itkin, T., Butler, J.M., et al. (2022). Engineering a niche supporting hematopoietic stem cell development using integrated single-cell transcriptomics. Nature communications 13, 1584. 10.1038/s41467-022-28781-z.

31. Tober, J., Maijenburg, M.M.W., Li, Y., Gao, L., Hadland, B.K., Gao, P., Minoura, K., Bernstein, I.D., Tan, K., and Speck, N.A. (2018). Maturation of hematopoietic stem cells from prehematopoietic stem cells is accompanied by up-regulation of PD-L1. J Exp Med 215, 645–659. 10.1084/jem.20161594.

32. Batsivari, A., Rybtsov, S., Souilhol, C., Binagui-Casas, A., Hills, D., Zhao, S., Travers, P., and Medvinsky, A. (2017). Understanding Hematopoietic Stem Cell Development through Functional Correlation of Their Proliferative Status with the Intra-aortic Cluster Architecture. Stem cell reports 8, 1549–1562. 10.1016/j.stemcr.2017.04.003.

33. Zhou, F., Li, X., Wang, W., Zhu, P., Zhou, J., He, W., Ding, M., Xiong, F., Zheng, X., Li, Z., et al. (2016). Tracing haematopoietic stem cell formation at single-cell resolution. Nature 533, 487–492. 10.1038/nature17997.

34. Li, Y., Gao, L., Hadland, B., Tan, K., and Speck, N.A. (2017). CD27 marks murine embryonic hematopoietic stem cells and type II prehematopoietic stem cells. Blood 130, 372–376. 10.1182/blood-2017-03-776849.

35. de Ceuninck van Capelle, C., Spit, M., and Ten Dijke, P. (2020). Current perspectives on inhibitory SMAD7 in health and disease. Crit Rev Biochem Mol Biol 55, 691–715. 10.1080/10409238.2020.1828260.

36. Monteiro, R., Pinheiro, P., Joseph, N., Peterkin, T., Koth, J., Repapi, E., Bonkhofer, F., Kirmizitas, A., and Patient, R. (2016). Transforming Growth Factor beta Drives Hemogenic Endothelium Programming and the Transition to Hematopoietic Stem Cells. Developmental cell 38, 358–370. 10.1016/j.devcel.2016.06.024.

37. Lempereur, A., Canto, P.Y., Richard, C., Martin, S., Thalgott, J., Raymond, K., Lebrin, F., Drevon, C., and Jaffredo, T. (2018). The TGFbeta pathway is a key player for the endothelial-to-hematopoietic transition in the embryonic aorta. Dev Biol 434, 292–303. 10.1016/j.ydbio.2017.12.006.

38. Ottersbach, K. (2019). Endothelial-to-haematopoietic transition: an update on the process of making blood. Biochem Soc Trans 47, 591–601. 10.1042/BST20180320.

39. Howell, E.D., Yzaguirre, A.D., Gao, P., Lis, R., He, B., Lakadamyali, M., Rafii, S., Tan, K., and Speck, N.A. (2021). Efficient hemogenic endothelial cell specification by RUNX1 is dependent on baseline chromatin accessibility of RUNX1-regulated TGFbeta target genes. Genes Dev 35, 1475–1489. 10.1101/gad.348738.121.

40. Sorensen, I., Adams, R.H., and Gossler, A. (2009). DLL1-mediated Notch activation regulates endothelial identity in mouse fetal arteries. Blood 113, 5680–5688. blood-2008-08-174508 [pii] 10.1182/blood-2008-08-174508.

41. Yokomizo, T., and Dzierzak, E. (2010). Three-dimensional cartography of hematopoietic clusters in the vasculature of whole mouse embryos. Development 137, 3651–3661. dev.051094 [pii] 10.1242/dev.051094.

42. North, T.E., de Bruijn, M.F., Stacy, T., Talebian, L., Lind, E., Robin, C., Binder, M., Dzierzak, E., and Speck, N.A. (2002). Runx1 expression marks long-term repopulating hematopoietic stem cells in the midgestation mouse embryo. Immunity 16, 661–672. S1074761302002960 [pii].

43. Tober, J., Yzaguirre, A.D., Piwarzyk, E., and Speck, N.A. (2013). Distinct temporal requirements for Runx1 in hematopoietic progenitors and stem cells. Development 140, 3765–3776. 10.1242/dev.094961.

44. Boisset, J.C., Clapes, T., Klaus, A., Papazian, N., Onderwater, J., Mommaas-Kienhuis, M., Cupedo, T., and Robin, C. (2015). Progressive maturation toward hematopoietic stem cells in the mouse embryo aorta. Blood 125, 465–469. 10.1182/blood-2014-07-588954.

45. Masuda, K., Kubagawa, H., Ikawa, T., Chen, C.C., Kakugawa, K., Hattori, M., Kageyama, R., Cooper, M.D., Minato, N., Katsura, Y., and Kawamoto, H. (2005). Prethymic T-cell development defined by the expression of paired immunoglobulin-like receptors. Embo J 24, 4052–4060.

46. Berthault, C., Ramond, C., Burlen-Defranoux, O., Soubigou, G., Chea, S., Golub, R., Pereira, P., Vieira, P., and Cumano, A. (2017). Asynchronous lineage priming determines commitment to T cell and B cell lineages in fetal liver. Nat Immunol 18, 1139–1149. 10.1038/ni.3820.

47. Hadland, B.K., Varnum-Finney, B., Poulos, M.G., Moon, R.T., Butler, J.M., Rafii, S., and Bernstein, I.D. (2015). Endothelium and NOTCH specify and amplify aorta-gonad-mesonephros-derived hematopoietic stem cells. J Clin Invest 125, 2032–2045. 10.1172/JCI80137.

48. Hadland, B.K., Varnum-Finney, B., Nourigat-Mckay, C., Flowers, D., and Bernstein, I.D. (2018). Clonal Analysis of Embryonic Hematopoietic Stem Cell Precursors Using Single Cell Index Sorting Combined with Endothelial Cell Niche Co-culture. J Vis Exp. 10.3791/56973.

49. Ventura, A., Kirsch, D.G., McLaughlin, M.E., Tuveson, D.A., Grimm, J., Lintault, L., Newman, J., Reczek, E.E., Weissleder, R., and Jacks, T. (2007). Restoration of p53 function leads to tumour regression in vivo. Nature 445, 661–665. 10.1038/nature05541.

50. Morino-Koga, S., and Yokomizo, T. (2024). Deciphering hematopoietic stem cell development: key signaling pathways and mechanisms. Front Cell Dev Biol 12, 1510198. 10.3389/fcell.2024.1510198.

51. Baron, C.S., Kester, L., Klaus, A., Boisset, J.C., Thambyrajah, R., Yvernogeau, L., Kouskoff, V., Lacaud, G., van Oudenaarden, A., and Robin, C. (2018). Single-cell transcriptomics reveal the dynamic of haematopoietic stem cell production in the aorta. Nature communications 9, 2517. 10.1038/s41467-018-04893-3.

52. Fadlullah, M.Z.H., Neo, W.H., Lie, A.L.M., Thambyrajah, R., Patel, R., Mevel, R., Aksoy, I., Do Khoa, N., Savatier, P., Fontenille, L., et al. (2022). Murine AGM single-cell profiling identifies a continuum of hemogenic endothelium differentiation marked by ACE. Blood 139, 343–356. 10.1182/blood.2020007885.

53. Souilhol, C., Gonneau, C., Lendinez, J.G., Batsivari, A., Rybtsov, S., Wilson, H., Morgado-Palacin, L., Hills, D., Taoudi, S., Antonchuk, J., et al. (2016). Inductive interactions mediated by interplay of asymmetric signalling underlie development of adult haematopoietic stem cells. Nature communications 7, 10784. 10.1038/ncomms10784.

54. Thambyrajah, R., and Bigas, A. (2022). Notch Signaling in HSC Emergence: When, Why and How. Cells 11. 10.3390/cells11030358.

55. Richard, C., Drevon, C., Canto, P.Y., Villain, G., Bollerot, K., Lempereur, A., Teillet, M.A., Vincent, C., Rossello Castillo, C., Torres, M., et al. (2013). Endothelio-mesenchymal interaction controls runx1 expression and modulates the notch pathway to initiate aortic hematopoiesis. Developmental cell 24, 600–611. 10.1016/j.devcel.2013.02.011.

56. Lizama, C.O., Hawkins, J.S., Schmitt, C.E., Bos, F.L., Zape, J.P., Cautivo, K.M., Borges Pinto, H., Rhyner, A.M., Yu, H., Donohoe, M.E., et al. (2015). Repression of arterial genes in hemogenic endothelium is sufficient for haematopoietic fate acquisition. Nature communications 6, 7739. 10.1038/ncomms8739.

57. Porcheri, C., Golan, O., Calero-Nieto, F.J., Thambyrajah, R., Ruiz-Herguido, C., Wang, X., Catto, F., Guillen, Y., Sinha, R., Gonzalez, J., et al. (2020). Notch ligand Dll4 impairs cell recruitment to aortic clusters and limits blood stem cell generation. EMBO J 39, e104270. 10.15252/embj.2019104270.

58. Souilhol, C., Lendinez, J.G., Rybtsov, S., Murphy, F., Wilson, H., Hills, D., Batsivari, A., Binagui-Casas, A., McGarvey, A.C., MacDonald, H.R., et al. (2016). Developing HSCs become Notch independent by the end of maturation in the AGM region. Blood 128, 1567–1577. 10.1182/blood-2016-03-708164.

59. Thambyrajah, R., Maqueda, M., Neo, W.H., Imbach, K., Guillen, Y., Grases, D., Fadlullah, Z., Gambera, S., Matteini, F., Wang, X., et al. (2024). Cis inhibition of NOTCH1 through JAGGED1 sustains embryonic hematopoietic stem cell fate. Nature communications 15, 1604. 10.1038/s41467-024-45716-y.

60. Clements, W.K., and Khoury, H. (2024). The molecular and cellular hematopoietic stem cell specification niche. Exp Hematol 136, 104280. 10.1016/j.exphem.2024.104280.

61. Lawson, N.D., Scheer, N., Pham, V.N., Kim, C.H., Chitnis, A.B., Campos-Ortega, J.A., and Weinstein, B.M. (2001). Notch signaling is required for arterial-venous differentiation during embryonic vascular development. Development 128, 3675–3683.

62. Duarte, A., Hirashima, M., Benedito, R., Trindade, A., Diniz, P., Bekman, E., Costa, L., Henrique, D., and Rossant, J. (2004). Dosage-sensitive requirement for mouse Dll4 in artery development. Genes Dev 18, 2474–2478.

63. Krebs, L.T., Shutter, J.R., Tanigaki, K., Honjo, T., Stark, K.L., and Gridley, T. (2004). Haploinsufficient lethality and formation of arteriovenous malformations in Notch pathway mutants. Genes Dev 18, 2469–2473.

64. Gale, N.W., Dominguez, M.G., Noguera, I., Pan, L., Hughes, V., Valenzuela, D.M., Murphy, A.J., Adams, N.C., Lin, H.C., Holash, J., et al. (2004). Haploinsufficiency of delta-like 4 ligand results in embryonic lethality due to major defects in arterial and vascular development. Proc Natl Acad Sci U S A 101, 15949–15954.

65. Gama-Norton, L., Ferrando, E., Ruiz-Herguido, C., Liu, Z., Guiu, J., Islam, A.B., Lee, S.U., Yan, M., Guidos, C.J., Lopez-Bigas, N., et al. (2015). Notch signal strength controls cell fate in the haemogenic endothelium. Nature communications 6, 8510. 10.1038/ncomms9510.

66. Robert-Moreno, A., Guiu, J., Ruiz-Herguido, C., Lopez, M.E., Ingles-Esteve, J., Riera, L., Tipping, A., Enver, T., Dzierzak, E., Gridley, T., et al. (2008). Impaired embryonic haematopoiesis yet normal arterial development in the absence of the Notch ligand Jagged1. Embo J 27, 1886–1895.

67. Ikeno, S., Nakano, N., Sano, K., Minowa, T., Sato, W., Akatsu, R., Sakata, N., Hanagata, N., Fujii, M., Itoh, F., and Itoh, S. (2019). PDZK1-interacting protein 1 (PDZK1IP1) traps Smad4 protein and suppresses transforming growth factor-beta (TGF-beta) signaling. The Journal of biological chemistry 294, 4966–4980. 10.1074/jbc.RA118.004153.

68. Blank, U., Karlsson, G., Moody, J.L., Utsugisawa, T., Magnusson, M., Singbrant, S., Larsson, J., and Karlsson, S. (2006). Smad7 promotes self-renewal of hematopoietic stem cells. Blood 108, 4246–4254. 10.1182/blood-2006-02-005611.

69. Ding, L., Saunders, T.L., Enikolopov, G., and Morrison, S.J. (2012). Endothelial and perivascular cells maintain haematopoietic stem cells. Nature 481, 457–462. 10.1038/nature10783.

70. Khan, J.A., Mendelson, A., Kunisaki, Y., Birbrair, A., Kou, Y., Arnal-Estape, A., Pinho, S., Ciero, P., Nakahara, F., Ma’ayan, A., et al. (2016). Fetal liver hematopoietic stem cell niches associate with portal vessels. Science 351, 176–180. 10.1126/science.aad0084.

71. Sitnicka, E., Ruscetti, F.W., Priestley, G.V., Wolf, N.S., and Bartelmez, S.H. (1996). Transforming growth factor beta 1 directly and reversibly inhibits the initial cell divisions of long-term repopulating hematopoietic stem cells. Blood 88, 82–88.

72. Fortunel, N.O., Hatzfeld, A., and Hatzfeld, J.A. (2000). Transforming growth factor-beta: pleiotropic role in the regulation of hematopoiesis. Blood 96, 2022–2036.

73. Yamazaki, S., Iwama, A., Takayanagi, S., Eto, K., Ema, H., and Nakauchi, H. (2009). TGF-beta as a candidate bone marrow niche signal to induce hematopoietic stem cell hibernation. Blood 113, 1250–1256. 10.1182/blood-2008-04-146480.

74. Wang, X., Dong, F., Zhang, S., Yang, W., Yu, W., Wang, Z., Zhang, S., Wang, J., Ma, S., Wu, P., et al. (2018). TGF-beta1 Negatively Regulates the Number and Function of Hematopoietic Stem Cells. Stem cell reports 11, 274–287. 10.1016/j.stemcr.2018.05.017.

75. Warsi, S., Blank, U., Dahl, M., Hooi Min Grahn, T., Schmiderer, L., Andradottir, S., and Karlsson, S. (2021). BMP signaling is required for postnatal murine hematopoietic stem cell self-renewal. Haematologica 106, 2203–2214. 10.3324/haematol.2019.236125.

76. Ganuza, M., Hall, T., Myers, J., Nevitt, C., Sanchez-Lanzas, R., Chabot, A., Ding, J., Kooienga, E., Caprio, C., Finkelstein, D., et al. (2022). Murine foetal liver supports limited detectable expansion of life-long haematopoietic progenitors. Nat Cell Biol 24, 1475–1486. 10.1038/s41556-022-00999-5.

77. Ng, E.S., Sarila, G., Li, J.Y., Edirisinghe, H.S., Saxena, R., Sun, S., Bruveris, F.F., Labonne, T., Sleebs, N., Maytum, A., et al. (2024). Long-term engrafting multilineage hematopoietic cells differentiated from human induced pluripotent stem cells. Nat Biotechnol. 10.1038/s41587-024-02360-7.

78. Yokomizo, T., Yamada-Inagawa, T., Yzaguirre, A.D., Chen, M.J., Speck, N.A., and Dzierzak, E. (2012). Whole-mount three-dimensional imaging of internally localized immunostained cells within mouse embryos. Nat Protoc 7, 421–431. nprot.2011.441 [pii] 10.1038/nprot.2011.441.

79. Schindelin, J., Arganda-Carreras, I., Frise, E., Kaynig, V., Longair, M., Pietzsch, T., Preibisch, S., Rueden, C., Saalfeld, S., Schmid, B., et al. (2012). Fiji: an open-source platform for biological-image analysis. Nat Methods 9, 676–682. nmeth.2019 [pii] 10.1038/nmeth.2019.

80. Hu, Y., and Smyth, G.K. (2009). ELDA: extreme limiting dilution analysis for comparing depleted and enriched populations in stem cell and other assays. Journal of immunological methods 347, 70–78. 10.1016/j.jim.2009.06.008.

81. Kleiter, I., Song, J., Lukas, D., Hasan, M., Neumann, B., Croxford, A.L., Pedre, X., Hovelmeyer, N., Yogev, N., Mildner, A., et al. (2010). Smad7 in T cells drives T helper 1 responses in multiple sclerosis and experimental autoimmune encephalomyelitis. Brain 133, 1067–1081. 10.1093/brain/awq039.

82. Qiu, X., Mao, Q., Tang, Y., Wang, L., Chawla, R., Pliner, H.A., and Trapnell, C. (2017). Reversed graph embedding resolves complex single-cell trajectories. Nat Methods 14, 979–982. 10.1038/nmeth.4402.

83. Stuart, T., Butler, A., Hoffman, P., Hafemeister, C., Papalexi, E., Mauck, W.M., 3rd, Hao, Y., Stoeckius, M., Smibert, P., and Satija, R. (2019). Comprehensive Integration of Single-Cell Data. Cell 177, 1888-1902 e1821. 10.1016/j.cell.2019.05.031.

84. Chen, C., Yu, W., Alikarami, F., Qiu, Q., Chen, C.H., Flournoy, J., Gao, P., Uzun, Y., Fang, L., Davenport, J.W., et al. (2022). Single-cell multiomics reveals increased plasticity, resistant populations, and stem-cell-like blasts in KMT2A-rearranged leukemia. Blood 139, 2198–2211. 10.1182/blood.2021013442.

