## Supplemental Table 1 for "Restraint of TGFβ family signaling by SMAD7 is necessary for hematopoietic stem cell maturation in the embryo"

**Supplementary Table 2**

| Dataset/Cell Type | Surface Phenotype | # Embryos | Somite Pairs | # Cells Sequenced | Median # Expressed Genes | Median # UMIs |
| --- | --- | --- | --- | --- | --- | --- |
| E11.5 Smad7^f/f^ E+HE+IAHC | CD41^mid/lo^CD31^+^Kit^+^CD144^+^  ESAM^+^Ter119^-^ | 7 | 39-44 | 4055 | 7321 | 42686 |
| E11.5 Smad7^Δ/Δ^ E+HE+IAHC | CD41^mid/lo^CD31^+^Kit^+^CD144^+^  ESAM^+^Ter119^-^ | 8 | 39-44 | 4326 | 8044 | 58705 |
| E18.5 Smad7^f/f^  LT-HSCs | Kit^+^Sca-1^+^CD48^-^CD150^+^Ter119^-^B220^-^CD3e^-^Gr1^-^Nk1.1^-^ | 5 | Theiler Stage 26 | 1463 | 4178 | 14683 |
| E18.5 Smad7^Δ/Δ^  LT-HSCs | Kit^+^Sca-1^+^CD48^-^CD150^+^Ter119^-^B220^-^CD3e^-^Gr1^-^Nk1.1^-^ | 4 | Theiler Stage 26 | 2126 | 4103 | 15044 |
| E18.5 Smad7^f/f^  LK cells | Kit^+^Sca-1^-^CD48^-^CD150^+^Ter119^-^B220^-^CD3e^-^Gr1^-^Nk1.1^-^ | 5 | Theiler Stage 26 | 4964 | 4417 | 20138 |
| E18.5 Smad7^Δ/Δ^  LK Cells | Kit^+^Sca-1^-^CD48^-^CD150^+^Ter119^-^B220^-^CD3e^-^Gr1^-^Nk1.1^-^ | 4 | Theiler Stage 26 | 4363 | 4378 | 17878 |

**Supplementary Table 3**
