## Supplemental Table 2 for "Restraint of TGFβ family signaling by SMAD7 is necessary for hematopoietic stem cell maturation in the embryo"

**Supplemental Table 1. Complete List of Antibodies.**

| **Antibody** | **Clone** | **Fluorophore** | **Supplier** | **Application** | **RRID #** |
| --- | --- | --- | --- | --- | --- |
| c-Kit | 2B8 | Alexa Fluor 700 | Thermo Fisher | Flow | AB_657583 |
| c-Kit | 2B8 | APC- eFluor 780 | Thermo Fisher | Flow | AB_1272213 |
| c-Kit | 2B8 | FITC | Biolegend | Flow | AB_313215 |
| c-Kit | 2B8 | Unconjugated |  | Whole Mount |  |
| CD3e | 145-2C11 | APC | Biolegend | Flow | AB_312677 |
| CD3e | 145-2C11 | PE | Thermo Fisher | Flow | AB_465496 |
| CD19 | eBio1D3 (1D3) | APC | Thermo Fisher | Flow | AB_1659676 |
| CD31 (PECAM-1) | 390 | PE-Cy7 | Thermo Fisher | Flow | AB_2716949 |
| CD31 (PECAM-1) |  | Unconjugated |  | Whole Mount |  |
| CD41a | eBioMWReg30 (MWReg30) | PerCP-eFluor710 | Thermo Fisher | Flow | AB_10855042 |
| CD45 | 30-F11 | PerCP-Cy5.5 | Thermo Fisher | Flow | AB_906233 |
| CD45.1 | A20 | APC-Cy7 | Biolegend | Flow | AB_313505 |
| CD45.1 | A20 | PE-Cy7 | Thermo Fisher | Flow | AB_469629 |
| CD45.2 | 104 | FITC | Thermo Fisher | Flow | AB_465062 |
| CD45R/B220 | RA3-6B2 | APC | Biolegend | Flow | AB_312997 |
| CD48 | HM48-1 | eFluor450 | Thermo Fisher | Flow | AB_11151336 |
| CD144 | 11D4.1 | PE | BD Biosciences | Flow | AB_2737609 |
| CD150 | TC15-12F12.2 | PE-Cy7 | Biolegend | Flow | AB_439797 |
| CD201 | eBio1560 (1560) | APC | Thermo Fisher | Flow | AB_10717805 |
| Cytokeratin 8 | TROMA-1 | Unconjugated | DSHB | Whole Mount |  |
| ESAM | 1G8/ESAM | FITC | Biolegend | Flow | AB_2044017 |
| F4/80 | BM8 | APC-Cy7 | Biolegend | Flow | AB_893477 |
| Gr-1 | RB6-8C5 | APC | Biolegend | Flow | AB_313377 |
| Gr-1 | RB6-8C5 | APC-Cy7 | BD Biosciences | Flow | AB_396775 |
| Gr-1 | RB6-8C5 | PerCP-Cy5.5 | Thermo Fisher | Flow | AB_906247 |
| Mac-1 | M1/70 | APC | Biolegend | Flow | AB_312795 |
| Mac-1 | M1/70 | APC-Cy7 | BD Biosciences | Flow | AB_396772 |
| Nk1.1 | PK136 | APC | Biolegend | Flow | AB_313397 |
| Runx1/Runx2/ Runx3AML1 |  | Unconjugated | Abcam | Whole Mount |  |
| Sca-1 | D7 | FITC | Thermo Fisher | Flow | AB_465333 |
| Sca-1 | D7 | PerCP-Cy5.5 | Thermo Fisher | Flow | AB_914372 |
| Ter119 | TER-119 | APC | Biolegend | Flow | AB_313713 |
| Ter119 | TER-119 | eFluor450 | Thermo Fisher | Flow | AB_1518808 |
| Goat anti-rabbit 488 |  | Alexa Fluor 488 | Abcam | Whole Mount |  |
| Goat anti-rat 555 |  | Alexa Fluor 555 | Abcam | Whole Mount |  |
| Goat anti-rabbit 647 |  | Alexa Fluor 647 | Abcam | Whole Mount |  |
