## Supplementary Figures for "Restraint of TGFβ family signaling by SMAD7 is necessary for hematopoietic stem cell maturation in the embryo"

### Supplementary Fig. 1

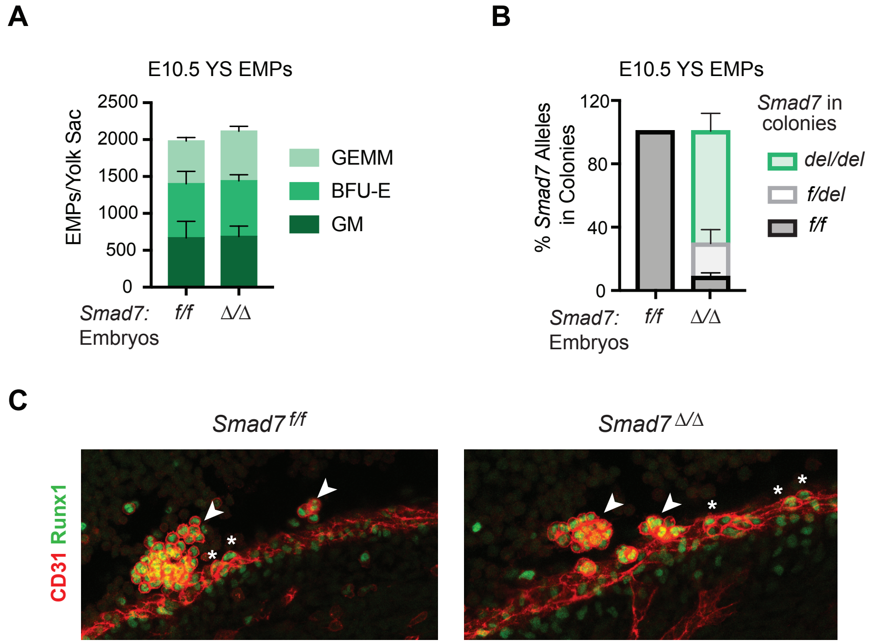

### Supplementary Fig. 2

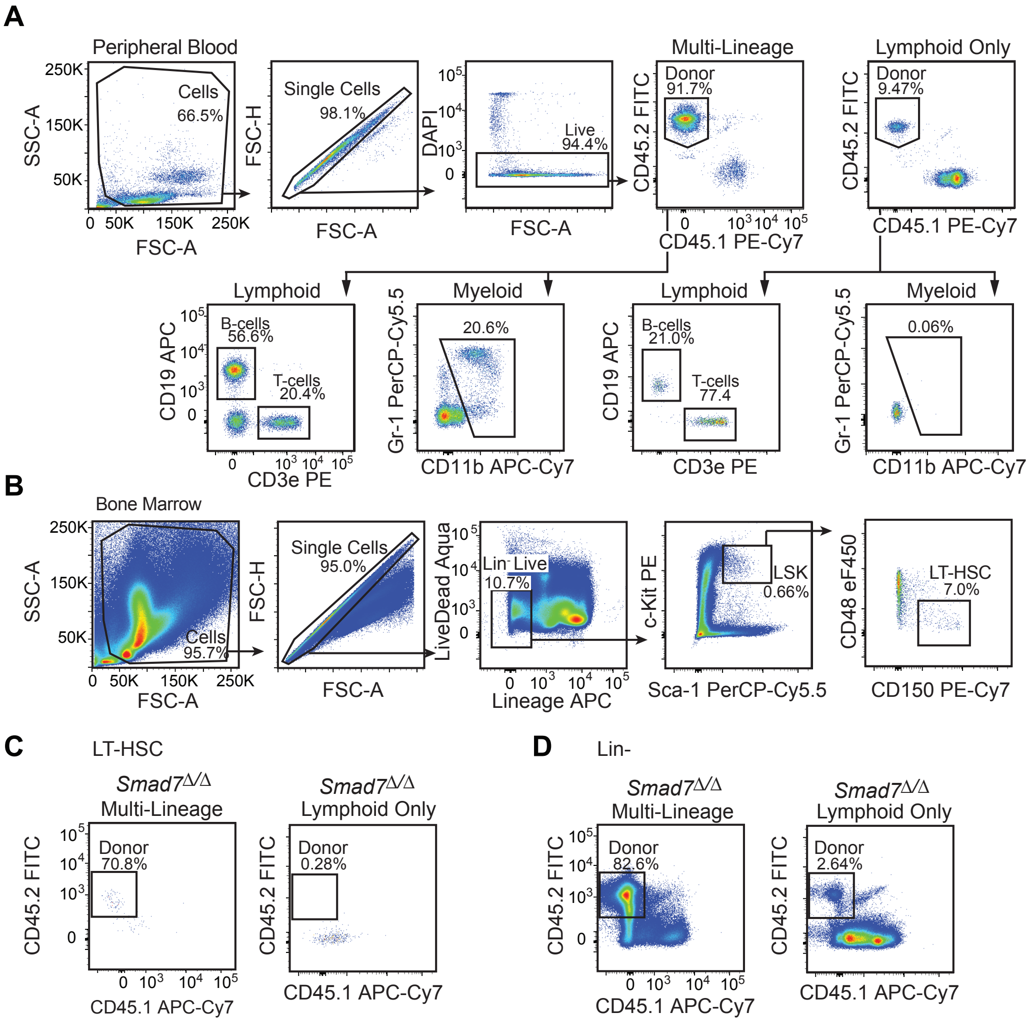

### Supplementary Figure 3

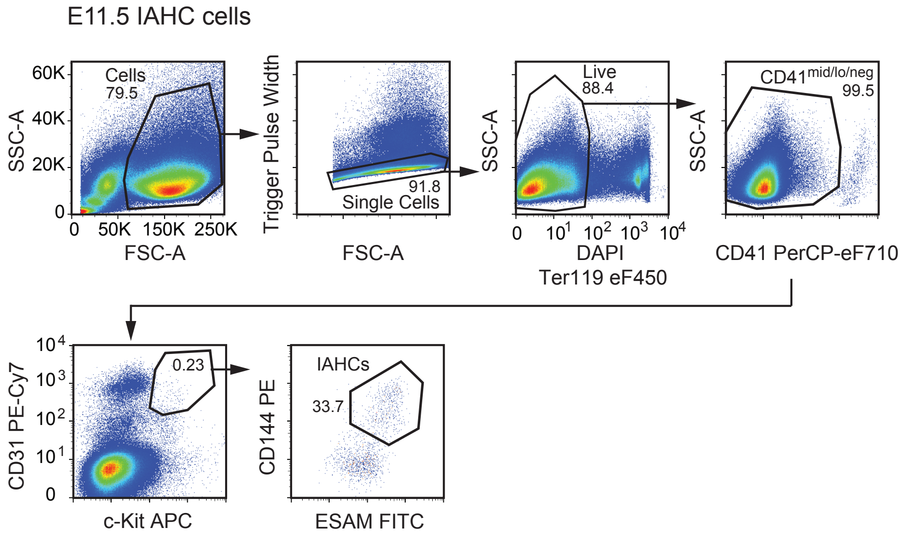

### Supplementary Fig. 4

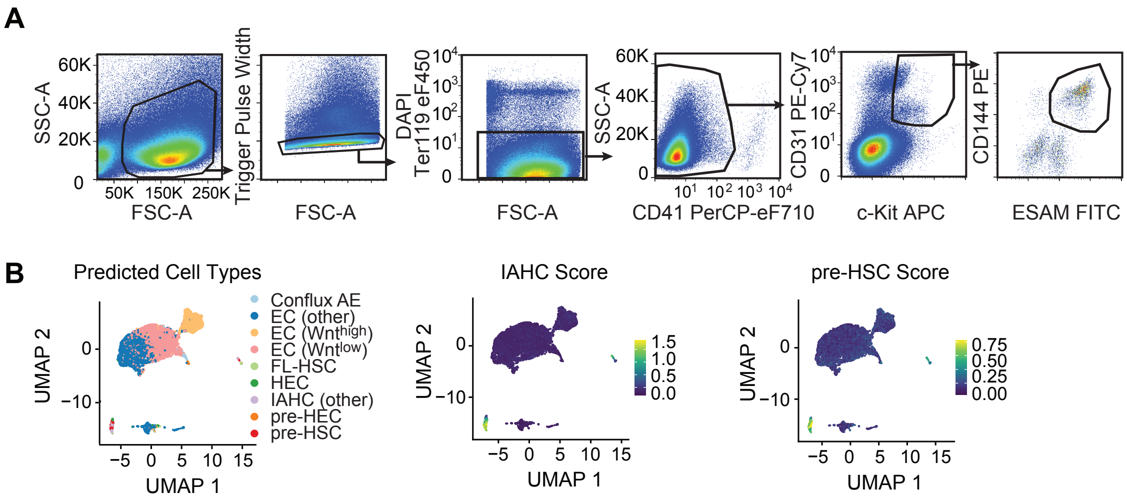

### Supplementary Fig. 5

A

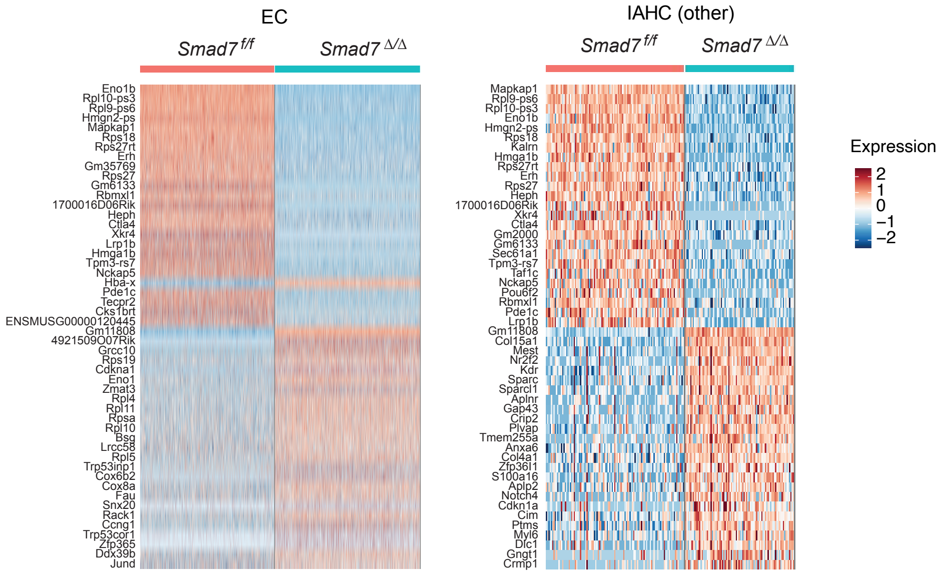


B

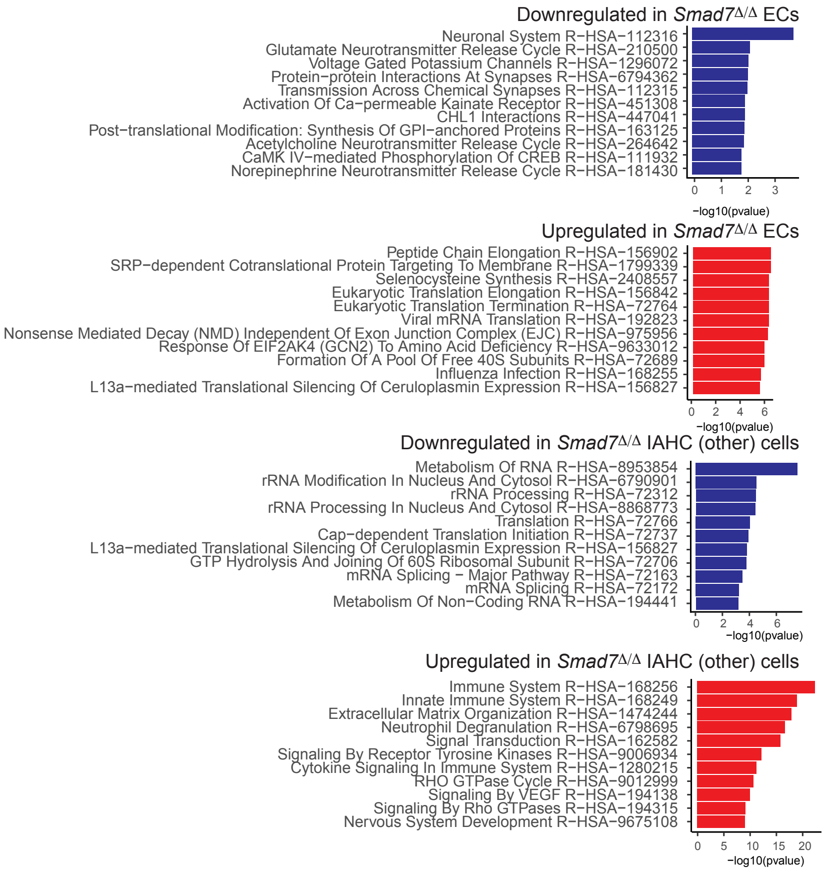

**Supplementary Fig. 6**

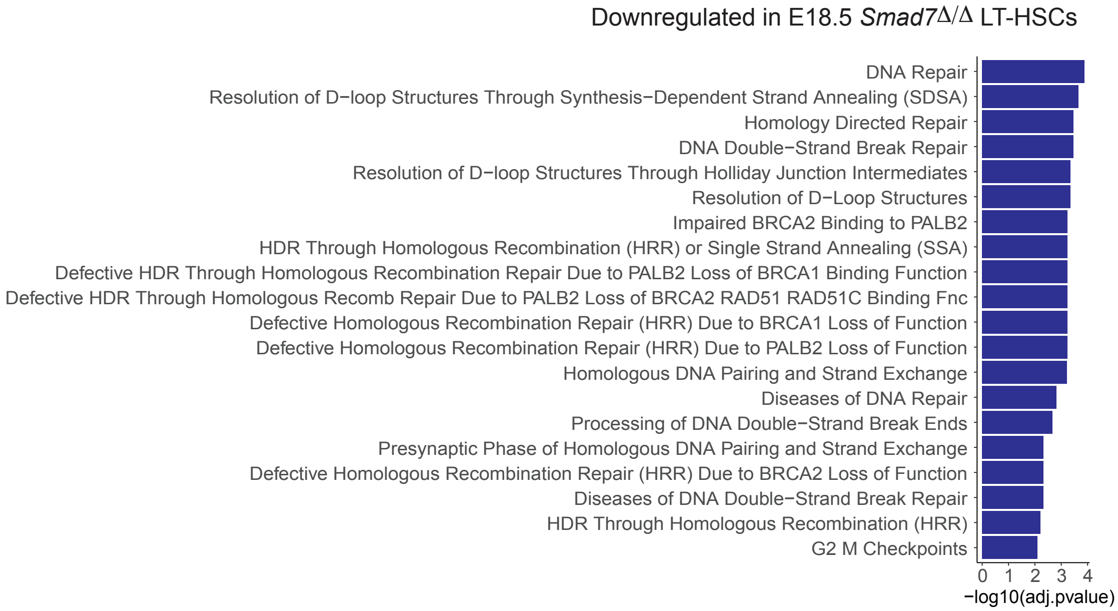
